# Filopodyan: an open-source pipeline for the analysis of filopodia

**DOI:** 10.1101/138610

**Authors:** Vasja Urbančič, Richard Butler, Benjamin Richier, Manuel Peter, Julia Mason, Frederick J. Livesey, Christine E. Holt, Jennifer L. Gallop

## Abstract

Filopodia have important sensory and mechanical roles in motile cells. The recruitment of actin regulators, such as ENA/VASP proteins, to sites of protrusion underlies diverse molecular mechanisms of filopodia formation and extension. We developed Filopodyan (<u>filopo</u>dia <u>dy</u>namics <u>an</u>alysis) in Fiji and R to measure fluorescence in filopodia and at their tips and bases concurrently with their morphological and dynamic properties. Filopodyan supports high-throughput phenotype characterization as well as detailed interactive editing of filopodia reconstructions through an intuitive graphical user interface. Our highly customizable pipeline is widely applicable, capable of detecting filopodia in four different cell types *in vitro* and *in vivo*. We use Filopodyan to quantify the recruitment of ENA and VASP preceding filopodia formation in neuronal growth cones, and uncover a molecular heterogeneity whereby different filopodia display markedly different responses to changes in the accumulation of ENA and VASP fluorescence in their tips over time.

**eTOC summary:** Urbančič et al. developed an open-source platform called Filopodyan (filopodia dynamics analysis) in Fiji and R to measure fluorescence in filopodia, at their tips, and bases, concurrently with their morphological and dynamic properties. This customizable tool therefore enables researchers to determine the relationship between protein localization and filopodium behavior.

## Introduction

Cells form an extensive network of actin-rich protrusions in order to move through tissues, including the veil and finger-like structures lamellipodia and filopodia (Bisi et al., 2013; Jacquemet et al., 2015; Krause and Gautreau, 2014; Mattila and Lappalainen, 2008). Filopodia play key roles throughout embryonic development, including in the developing nervous system. Filopodia mediate the sensing of attractant and repellent guidance cues by growth cones (McConnell et al., 2016; Zheng et al., 1996) and the guidance of axons through the developing nervous system (Bentley and Toroian-Raymond, 1986; Chien et al., 1993; O’Connor et al., 1990). Once neurites arrive at their destination, filopodia are involved in the formation of branches (Dwivedy et al., 2007) and synaptic connections (Lohmann and Bonhoeffer, 2008; Ziv and Smith, 1996).

Despite the overwhelming importance of filopodia, many questions about their biology remain unanswered. Firstly, the molecular mechanism of their formation is not clear. Multiple models have been proposed and experimentally supported, including (1) elongation of converging filaments within an underlying branched network of lamellipodial actin, fundamentally dependent on the Arp2/3 complex (Svitkina et al., 2003), (2) *de novo* nucleation of filopodial filaments at their tip by proteins also capable of promoting actin polymerization, such as formins (Faix and Rottner, 2006), and (3) initial clustering of membrane binding proteins which subsequently recruit other factors required for filopodium formation (Lee et al., 2010; Saarikangas et al., 2015). However, the molecular identities of protein assemblies driving filopodium formation according to these models are not clear-cut. ENA/VASP proteins localize to filopodia tips upon their protrusion and are implicated in regulating the number and length of filopodia in a variety of cell types, through diverse mechanisms that structurally fit the ‘convergent elongation’ and ‘tip nucleation’ models (Barzik et al., 2014; Lanier et al., 1999; Svitkina et al., 2003). In retinal ganglion cells, which we use here, ENA/VASP proteins are important for growth and stabilization of filopodia and terminal arborization of the axon (Dwivedy et al., 2007).

Moreover it also appears that cells use multiple filopodial actin filament elongating proteins. Studies in *Drosophila* and fibroblasts suggest that filopodia driven by ENA/VASP proteins or by formins (Dia/mDia2) have distinct properties (Barzik et al., 2014; Bilancia et al., 2014; Nowotarski et al., 2014). To better understand which proteins actively contribute to filopodial growth in various contexts, and to differentiate between models of filopodia formation it would be critical to quantify the amount of proteins of interest within filopodia concurrently with their dynamic behaviour. This approach has been very fruitful in understanding the protrusion of lamellipodia and membrane blebs (Barry et al., 2015; Charras et al., 2006; Lee et al., 2015; Machacek et al., 2009; Martin et al., 2016). A comprehensive image analysis pipeline is essential for these goals.

Recently computational tools have been developed for automated segmentation and analysis of filopodia. These include MATLAB applications FiloDetect, which measures the number and lengths of filopodia in non-neuronal cells (Nilufar et al., 2013), and CellGeo, which allows identification and tracking of filopodia and the assessment of phenotypes in their morphodynamic properties, as well as characterizing lamellipodial dynamics (Tsygankov et al., 2014). The ImageJ plugin ADAPT identifies filopodia, among its suite of other functions for automated quantification of cell migration and the dynamic behavior of lamellipodia and blebs in relation to their fluorescence intensities (Barry et al., 2015). Software has also been specifically designed for the automated detection of filopodia in dendrities, with a focus on their longitudinal and lateral movement (Hendricusdottir and Bergmann, 2014; Tarnok et al., 2015). Galic and colleagues (Saha et al., 2016) pioneered concurrent analysis of fluorescence and filopodium behaviour, developing MATLAB software for semi-automated analysis of filopodia dynamics and fluorescence. Most recently, the ImageJ plugin FiloQuant has enabled customizable automated quantification of filopodia lengths and densities, particularly in 3D microenvironments (Jacquemet et al., 2017, *Preprint*). All together, these studies demonstrate the surging demand for automated quantitative approaches in the field.

We built on these approaches, creating an ImageJ plugin for characterization of filopodial dynamics in parallel with the quantification of fluorescence, enabling us to determine the relationship between protein localization and filopodium behavior. Our plugin, Filopodyan, analyses fluorescence microscopy timelapse datasets using the open-source platforms FIJI (Schindelin et al., 2012) and R (R Core Team, 2016). We have used Filopodyan to track the recruitment of ENA and VASP to newly forming filopodia and to filopodia tips during their extension and retraction. By quantifying the cross-correlation between fluorescence intensity and tip movement, we find several subpopulations of filopodia with different molecular, though similar morphological behavior.

## Results

### A customizable pipeline for automated identification of filopodia in timelapse datasets

To allow the segmentation of filopodia within an image from the rest of the cell, we express a membrane marker within the cells of interest (Fig. 1 A). A membrane marker is more suitable for this purpose than a volume marker due to the high proportion of membrane to volume within filopodia relative to the rest of the cell, and we found that the uniform signal provided by a membrane marker performed better for reconstructions than an actin label, for example, SiR-actin. The contrast between signal and background at the edges of the cell is further amplified by applying a Laplacian-of-Gaussian (“LoG”) filter to the image. This enhances the outline of the cell boundary and a thresholding method is then applied to create a mask of total cell area. The scale of the LoG filter (σ) and the choice of thresholding method are set by the user (initial guidelines for choice of parameters depending on pixel dimensions are provided in the User Guide). A graphical user interface window enables a rapid preview of segmentation, assisting the user to identify most suitable segmentation parameters before processing the entire image stack. Examples of different parameters and the suitability of a range of values are shown in Fig. 1 B. We also included adaptive thresholding as an option that may be useful for reconstructing low signal-to-noise images. Filopodyan was developed using images taken with a 100x objective using a CMOS camera with 65 nm pixel width. It is possible to segment filopodia at bigger pixel dimensions and consequently a smaller number of pixels per filopodium width, but the reliability and accuracy of segmentation are reduced, such that segmentation is problematic when several filopodia are near each other.

**Figure 1.**
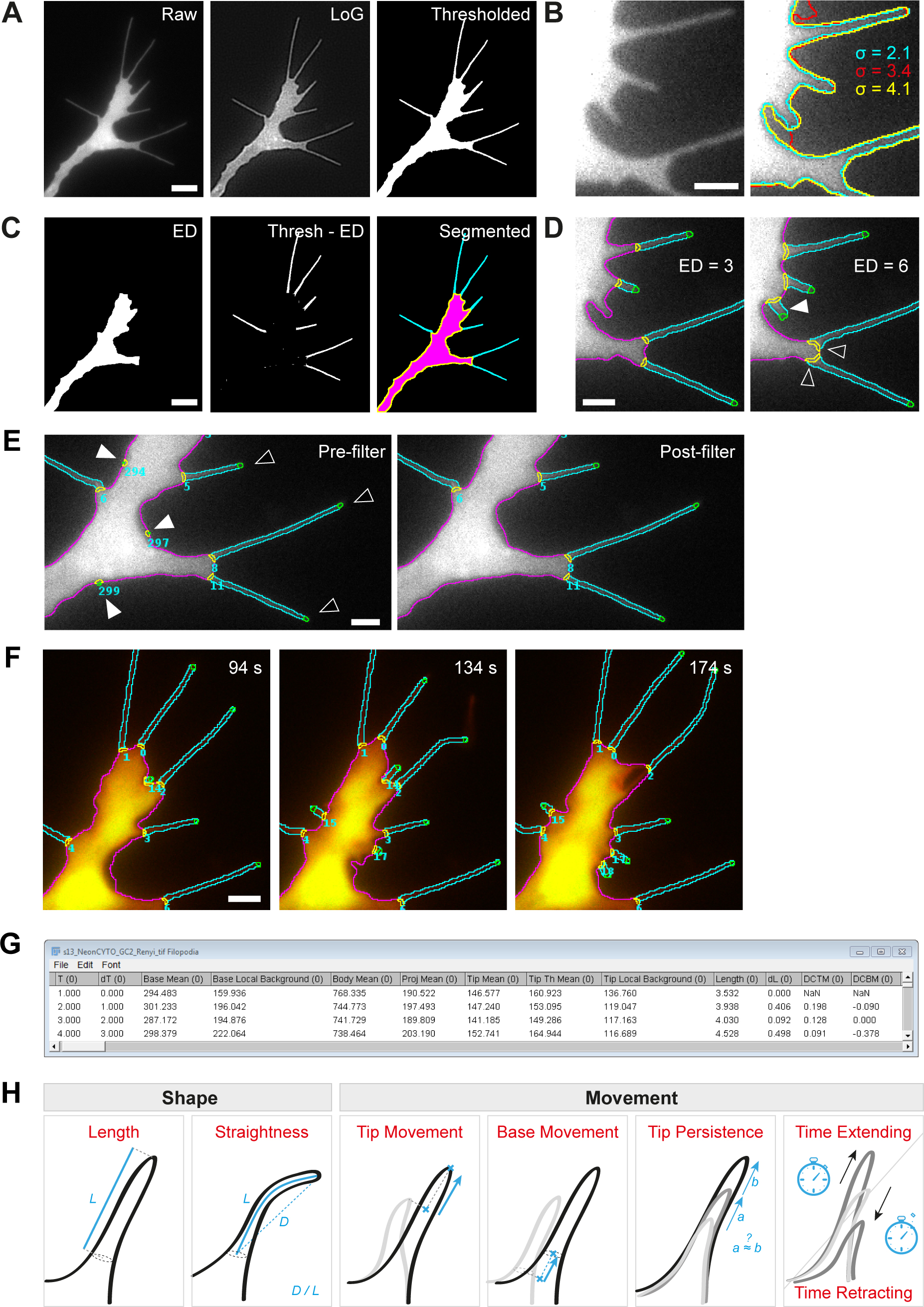
Filopodyan, a highly customizable pipeline for detection, tracking and analysis of protrusions in timelapse datasets. **(A)** Image of a *Xenopus* retinal ganglion cell growth cone with filopodia (left) is processed with a Laplacian of Gaussian (LoG) filter for edge contrast enhancement (middle), and binarised with a chosen threshold method (right). **(B)** Parameters for boundary mapping are customizable; coloured lines show the different cell boundaries detected by using three different values of LoG σ with the Renyi entropy thresholding method. **(C)** Iterative erosion-dilation (ED) of the thresholded mask (from A, right panel) removes protrusions (left), and the difference between these two masks yields a mask with protrusions alone (centre), leading to a fully segmented image (right; protrusions in cyan, growth cone body in magenta and its boundary in yellow). **(D)** The cutoff threshold for segmenting protrusions is fully user-customisable by adjusting the number of erosion-dilation operations. The two panels illustrate the difference upon changing the ED number from 3 to 6, showing additional protrusions (white arrowhead) and shifted position of their bases (black arrowheads). **(E)** Automated filtering removes false hits that do not satisfy user-defined selection criteria. Detected candidate protrusions include true (black arrowheads) as well as erroneous (white arrowheads) annotations which are removed by the filter. Any false hits that escape automated filtering and incorrect tracking can be corrected manually. **(F)** A time series of images segmented into body (magenta), protrusions (cyan), their bases (orange) and tips (green). **(G)** Output tables contain the measurements at each timepoint (row) of protrusion shape, movement, fluorescence and coordinates (column) for all detected protrusions (column sections). **(H)** Descriptive properties of filopodia extracted by Filopodyan plugin and the downstream analysis scripts include length, straightness, direction-corrected measures of tip movement and base movement, tip persistence and the time spent extending, retracting or stalling. Scale bars: 5 μm (A, C), 2 μm (B, D, E, F)

Because filopodia are thin protrusions, eroding the image mask that demarcates the total cell area readily removes them from the total cell mask (Fig. 1 C). Subsequent dilation re-expands the size of the cell footprint, and the subtraction of this image from the original thresholded image leaves the mask of protrusions alone. The image can thus be partitioned into separate compartments of cell body and protrusions. This procedure is similar as that used by previously developed filopodia detection tools FiloDetect and ADAPT (Barry et al., 2015; Nilufar et al., 2013). The number of steps for erosion and dilation is user-adjustable as the optimal value is dependent on the morphological properties of the imaged cell type as well as imaging parameters such as pixel size and the purpose of the analysis (Fig. 1 D). Once protrusions are thus identified as structures separate from the cell body, it is necessary to assign their base and tip positions. This is required for tracking tip movement and base movement over time, and for measuring protein fluorescence at the point on the cell body where the filopodium first protrudes and at the tip of the filopodium once it has formed. Filopodyan assigns the base and tip positions by annotating the positions within the mask of each protrusion that are nearest or furthest away from the cell body.

In order to measure extension and retraction events, filopodia need to be tracked over time. Structure identity is evaluated on the basis of (1) the time elapsed between the two recorded structures, (2) the distance between their base coordinates, (3) the distance between their tip coordinates, and (4) the overlap between their area in the two timepoints. Once the identity of structures over time is established, the dynamic parameters for each tracked protrusion are calculated (e.g. change in length, tip movement and base movement etc).

### Error-correcting capabilities

Some of the candidate structures initially detected are true hits, whereas others represent patches of higher pixel intensity noise near the boundary of the cell (Fig. 1 E). To distinguish between true and false hits Filopodyan performs an automated filtering step on detected candidate structures. User-defined thresholds are used to eliminate candidates that do not satisfy required criteria, which include minimum number of frames in existence, the time of appearance, maximum length reached during the timelapse, maximum tip movement during the timelapse, and mean waviness. The values for these thresholds are customizable and need empirical adjustment for each application. A rapid preview visualizes which structures are retained with the currently set threshold values, helping the user to select useful thresholds for the filtering. A second round of automated tracking is then performed on filtered hits, yielding an automated reconstruction of tracked filopodia identities over time throughout the duration of the timelapse.

Errors in automated tracking can occur, for instance when filopodia move laterally, and in addition some false positives may persist after automated filtering. For this reason we included a module for manual editing of the automatically reconstructed tracks in our plugin workflow. The user can inspect the reconstruction at each timepoint and reject candidate structures in individual timepoints or across the entire timelapse. The user can also correct errors in automated identity tracking by reassigning the identity of structures or creating identity links between different tracks.

### Output parameters

Filopodyan outputs a fully segmented timelapse with separate regions of interest demarcating the body, protrusion, base and tip segments (Video 1, Fig. 1 F), and produces data tables containing the measured properties of each identified structure at each timepoint, as well as their coordinates and the corresponding fluorescence measurements (Fig. 1 G).

The directly measured morphodynamic parameters of filopodia in the Filopodyan data table are: length (calculated from the perimeter value, with corrections applied for the width at the base and curvature at the tip), change in length between successive timepoints, tip movement from previous timepoint (corrected for lateral filopodium movement) and base movement from previous timepoint (also corrected for lateral movement) (Fig. 1 H). Further filopodium properties are derived from these first-order parameters by downstream R scripts, including: the persistence (autocorrelation) of tip movement, the proportions of time that the tip spends extending, retracting or stalling, and similarly the proportion of time that the base spends invading, retracting or in a stable state. Thresholds for extension or retraction are adjustable according to the application; in our case, if mean tip movement over a time window of 10 s exceeded 32.5 nm/s (equivalent to one pixel width per timepoint at the imaging parameters we used), the tip was assigned to the extending state at that timepoint, and if it was below –32.5 nm/s, the tip was assigned to the retracting state. Equally, base movement was assigned to invading state if its mean movement over 10 s exceeded 32.5 nm/s and to retracting state if it was below –32.5 nm/s. Rolling mean (smoothing across five successive timepoints) was applied to the measurements of tip and base movement to reduce noise.

To verify the quality of segmentation, we compared filopodium lengths as determined by manual line tracing to lengths of the same structures as computed by Filopodyan (Fig. S1 A), revealing good correlation between the lengths of filopodia in the same timelapse measured manually or with Filopodyan (n = 186 measurements; Pearson R = 0.97).

### Robust filopodia segmentation, tracking and measurement across cell types

We developed Filopodyan for the growth cones in *Xenopus* retinal ganglion cell (RGC) axons (Fig. 2 A, Video 1), a model system where the important role of filopodia for guidance and branching has been previously characterized *in vivo* (Chien et al., 1993; Dwivedy et al., 2007). We have additionally tested Filopodyan in several cell types in the developing *Drosophila* embryo due to the importance of this genetic model system. A combination of automated detection with some manual editing enables accurate detection of filopodia in *Drosophila* tracheal cells and in epithelial border cells during dorsal closure *in vivo* (Fig. 2 B-C, Videos 2-3).

**Figure 2.**
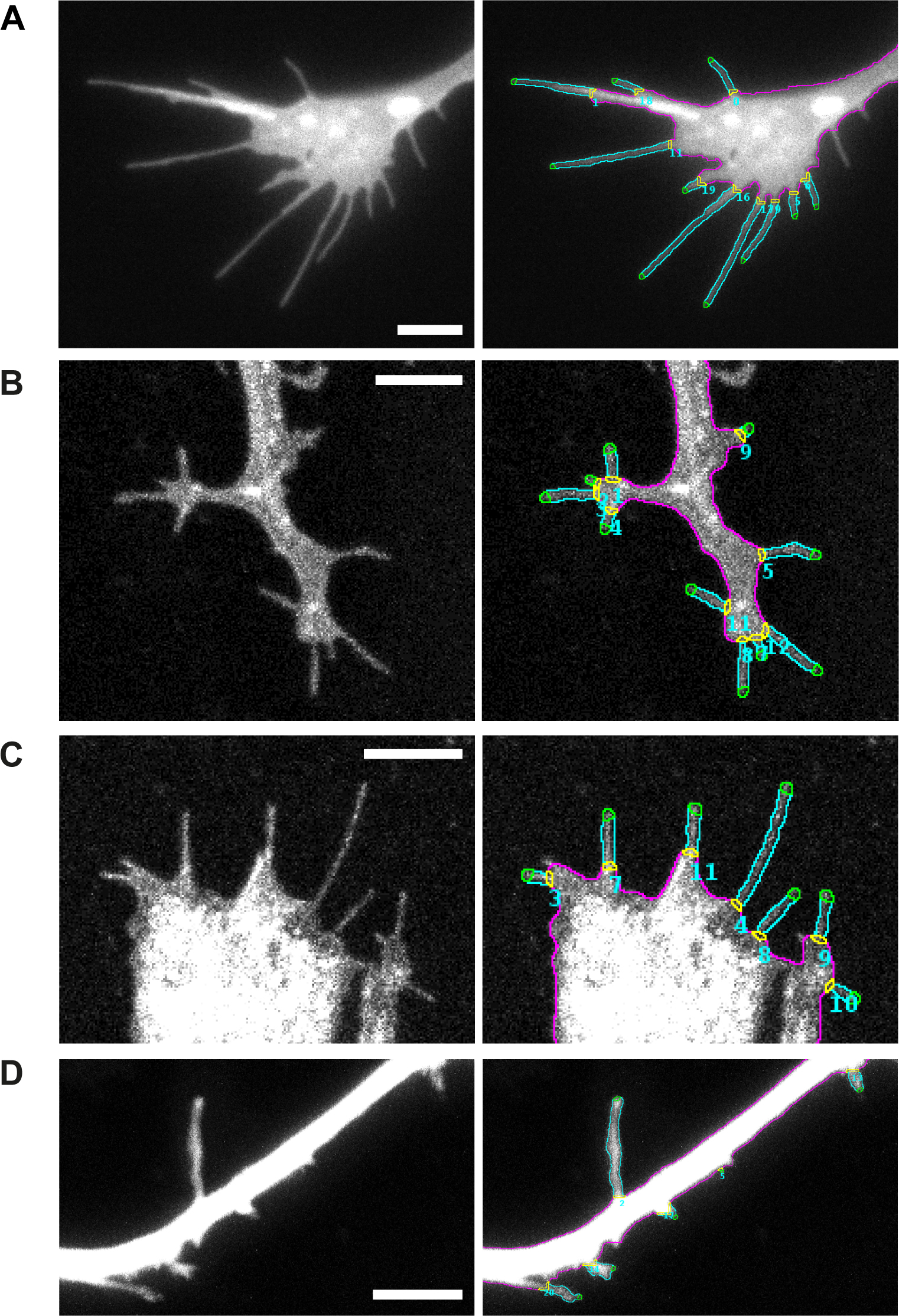
Filopodyan detects and accurately segments filopodia from a variety of different cell types *in vitro* and *in vivo*. **(A)** Growth cone from *Xenopus* retinal ganglion cell axons in culture, expressing GAP-RFP; single timepoint from Video 1. **(B)** Tracheal cells in a *Drosophila* embryo expressing *btl-Gal4 UAS-Cherry-CAAX*; a single timepoint from Video 2. **(C)** Leading edge cells during dorsal closure in a *Drosophila* embryo expressing *en-Gal4 UAS-cd8mCherry*; a single timepoint from Video 3. **(D)** Human iPS-cell derived cortical neurons in culture, expressing cytoplasmic mNeonGreen; a single timepoint from Video 4. Scale bars: 5 μm.

Induced pluripotent stem cell (iPSC)-derived human cortical neurons are an important model for studying neurodevelopmental and neurodegenerative conditions, including Down syndrome and Alzheimer’s disease (Shi et al., 2012). Dendrites of cortical neurons exhibit dynamic filopodia, which are precursors of dendritic spines (Hotulainen and Hoogenraad, 2010), and as such an important structure relevant for the development of stable synapses that underlie learning and memory (Xu et al., 2009; Yang et al., 2009). We imaged the dendrites of human iPSC-derived cortical neurons in culture using two-photon microscopy, and demonstrate that Filopodyan readily segments dendritic filopodia in stack projections (Fig. 2 D, Video 4). Exceptions to the versatility of Filopodyan are cell types with a very high density of filopodia and a high prevalence of filopodial crossing. In principle this can be resolved to some extent by removing selected filopodia in the manual editing step, but other software is likely to be more suitable in these cases. Other limitations of the software include its inability to handle branching and looping events.

### High-throughput extraction of filopodial features yields a content-rich dataset of filopodial dynamics from retinal ganglion cell growth cones

We used Filopodyan to generate a descriptive dataset of filopodial properties in *Xenopus* RGC growth cones. We analysed filopodia in timelapse videos of 19 growth cones, each video lasting 4 minutes and captured at the frame rate of 2 s/frame. We carefully curated the automated reconstructions by manually verifying each reconstructed filopodium at each timepoint and rejecting inaccurate segmentations (spurious detections of noise near the growth cone boundary, as well as filopodia branching, looping, or projecting from the axon shaft).

The properties of the manually curated dataset of 160 growth cone filopodia are summarized in Table S1 and Fig. 3 A. These measurements are similar to previously reported filopodia measurements. Filopodium lengths were 8.0 ± 4.8 μm (mean ± SD; maximum length during timelapse), and 5.6 ± 3.9 μm (mean length during timelapse), which compares to similar measurements for filopodia in *Xenopus* RGC growth cones *in vivo* (5.6 μm) (Atkinson-Leadbeater et al., 2016). Mean filopodium lengths of 7.8 μm and 7.2 μm have been reported in chick RGCs (Chen et al., 2006; Gehler et al., 2004) and 7.7 ± 3.6 μm in mouse dorsal root ganglion growth cones (McConnell et al., 2016). Our measured filopodial extension rate of 66 ± 26 nm/s (mean ± SD of the median extension rate per filopodium) is similar to previously reported rates in mouse superior cervical ganglion neurons (55 nm/s) (Brown and Bridgman, 2003), in dorsal root ganglion neurons (59 ± 3 nm/s) (McConnell et al., 2016) and in *Drosophila* primary neurons (87 nm/s) (Sanchez-Soriano et al., 2010).

**Figure 3.**
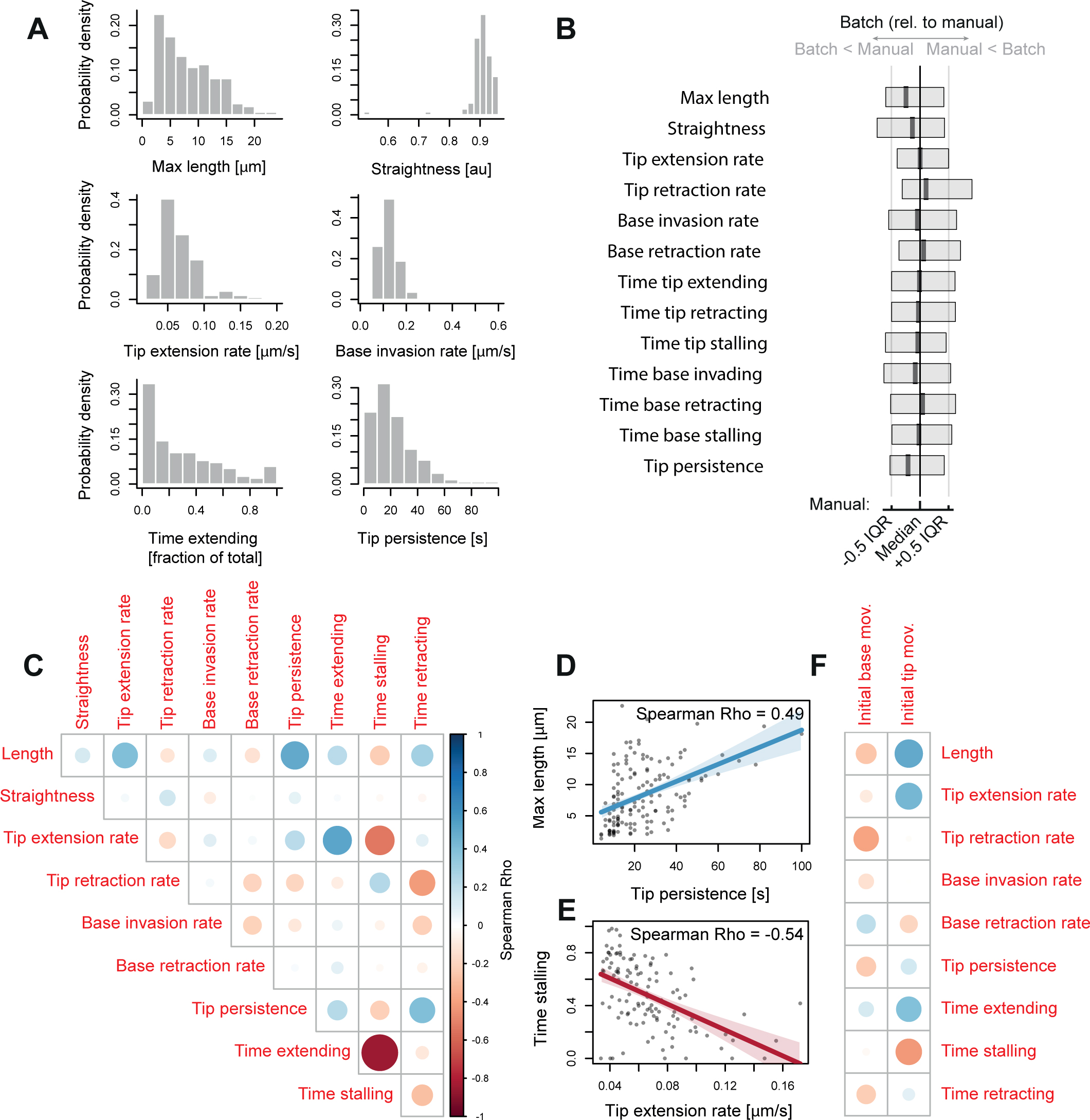
Correlations between parameters of filopodial dynamics. **(A)** Properties of *Xenopus* RGC growth cone filopodia from a manually curated dataset of (n = 160 filopodia, N = 19 neurons). Distribution histograms are shown for selected parameters: maximum length reached during timelapse, straightness at maximum length, median rate of tip extension during the timelapse, median rate of base invasion during the timelapse, proportion of time spent in extending state, and tip persistence; full description in Methods. Descriptive statistics for these and all other parameters are in Table S1. **(B)** Comparison of the filopodia phenotype between two analyses of the same dataset: the manually curated analysis and fully automated batch analysis. For each parameter, the normalized boxplot represents the difference of the batch analysis relative to the manual analysis; black and grey vertical lines represent the median and interquartile range of the manually curated dataset for each parameter. Boxes and thick vertical lines represent the interquartile ranges and medians of each parameter for the fully automated (batch) dataset. Full descriptive statistics are in Table S1. **(C)** Correlation matrix between parameters describing filopodium shape and movement in a dataset of *Xenopus* RGC growth cone filopodia (n = 160) expressing GAP-RFP and mNeonGreen. Colour scaling indicates the sign of correlation (blue-to-red for positive-to-negative), circle size indicates correlation strength. **(D)** Positive correlation between tip persistence of a filopodium and its maximum reached length. **(E)** Negative correlation between the median rate of tip extension of a filopodium and its proportion of time spent stalling. **(F)** Correlation matrix visualizing the correlations between the initial movement of the filopodial tips and bases (in the first 20 s after their initial appearance) with other filopodial properties in subsequent timepoints (after initial 20 s).

Manual curation is a time-intensive procedure and is prone to introducing subjective errors or human inconsistency. We thus tested how much the fidelity of reconstruction is preserved by relying only on the automated filtering step without human editing. This approach is potentially error prone as there is no mechanism to correct errors obvious to the human eye. However, when analyzing aggregate summary data from this dataset, we found that the results were mostly comparable to the results obtained after careful user curation (Fig. 3 B; Table S1; Fig. S1 B-I) – out of 13 parameters, the only parameter significantly different between the two analyses was filopodium length (6.6 ± 4.8 µm max length in batch mode, 8.0 ± 4.8 in manually curated dataset, mean ± SD, Holm-adjusted Mann-Whitney p = 0.011; Table S1, Fig. S1 B). By comparison to the manually curated dataset, which took 6 hours of user input to generate (verifying the accuracy of reconstruction for >160 structures in 121 timepoints), the fully automated dataset was generated in 15 min using batch mode with no further human input. Thus, when image quality is sufficiently high, Filopodyan is suitable for use in rapid high-throughput characterization and phenotyping of filopodial dynamics in large datasets.

Each filopodium is characterized by a set of values describing its dynamic behavior across the entire timelapse (maximum length reached, straightness at maximum length, median rate of extension while extending, time spent extending, etc.) – numbers that together provide a summary of its morphodynamic state throughout the timelapse. To gain new insights into the relationships between these parameters, we generated correlation matrices to mine for significant correlations (Fig. 3 C). The parameter that most strongly correlated with maximum length reached by a filopodium was its persistence of tip movement (Spearman Rho = 0.49, Holm-adjusted p = 1.24 ×10^-8^; Fig. 3 D). The rate of tip extension also positively correlated with maximum length (Rho = 0.40, p = 2.02 ×10^-4^); this agrees with the intuitive understanding that those filopodia that extend faster and whose tips display higher persistence of movement reach greater lengths. Also notably strong were the correlations indicating that filopodia with faster-extending tips spend more time extending (Rho = 0.52, p = 6.53 ×10^-8^) and less time stalling (Rho = −0.54, p = 1.11 ×10^-8^; Fig. 3 E).

Some filopodia originate by forward extension (tip-driven formation) whereas others are formed mostly as a result of lamellipodium retraction (base-driven formation), with a continuum of states between the extremes. We asked whether these formation mechanisms affected the subsequent properties later in their lifetimes. In our dataset of *Xenopus* retinal ganglion cell filopodia, the filopodia initiating with a greater tip extension rate during the first 20 s of their existence (‘initial tip movement’) reached longer maximum lengths (Fig. 3 F; Spearman Rho = 0.49, Holm-adjusted p-value = 0.0283), despite initial extension not strongly correlating with median tip extension rate later in their life (Rho = 0.44, p = 0.3). For initial base movement, no correlations with other parameters below the significance threshold of α = 0.05 were detected (Fig. 3 F).

We next asked whether filopodia that form without tip protrusion can later extend their tips. A third of filopodia in our dataset began their life with initial tip movement below the tip extension threshold (“non-protrusive” formation, 17/45 filopodia; Fig. S2 A). We plotted direction-corrected vector measures of tip movement, base movement and length against time, to reveal the how much the changes in filopodium length are a consequence of tip movement or base movement (Fig. S2 B, showing one example from protrusive and non-protrusive classes). As expected, protrusive filopodia later extend their tips, and furthermore non-protrusive filopodia were also able to sustain a tip extension later in their lifetime (Fig. S2 C). This observation is consistent with the idea that various finger-shaped protrusions (filopodia and retraction fibers) do not occupy rigid categories, but instead exist within dynamically interconvertible states (Svitkina et al., 2003). Non-protrusive filopodia attained lower length overall (Fig. S2 C, right panel), as expected from correlation between length and the rate of initial tip extension (Fig. 3 F).

### Predicting base position before filopodia formation

In order to address the question of the timing of recruitment of various proteins during filopodia formation, we sought to quantify fluorescence at initation sites on the membrane before projection of nascent filopodia. To do this we equipped Filopodyan with a feature to map predicted base positions of future filopodia in the timepoints preceding their formation: for each newly formed filopodium, Filopodyan maps the XYT coordinates of the point of origin (during the first timepoint of its existence), and in the timepoints preceding its formation the position of the future filopodium base is projected onto the edge-proximal area, thus measuring the fluorescence closest to the base of the future filopodium, to the best approximation (Fig. 4 A). This can be extended for a user-defined number of timepoints into the past from the moment of origin.

**Figure 4.**
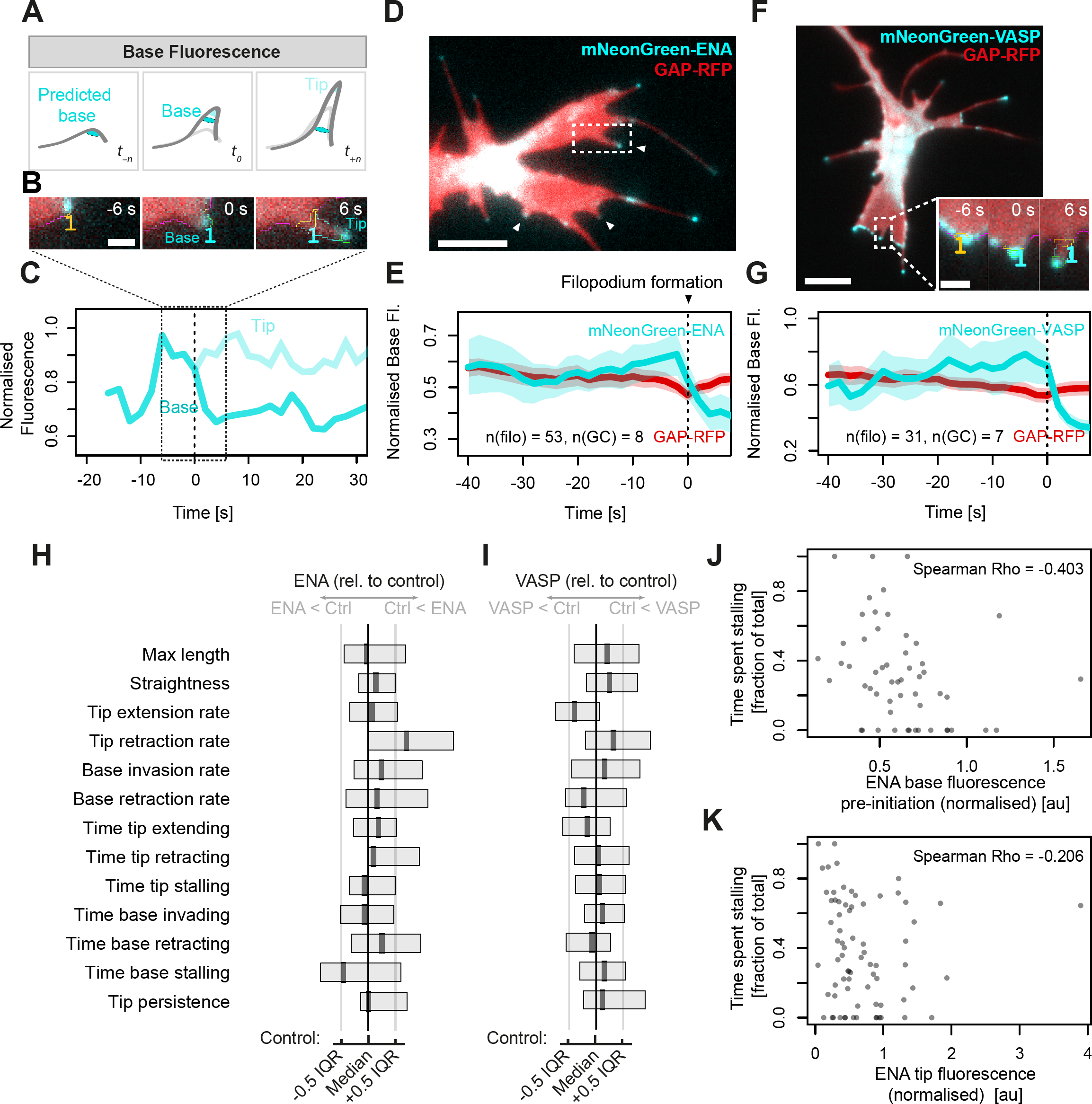
Predicted filopodium base position enables the measurement of protein recruitment before filopodium formation. **(A)** Measurements of fluorescence are taken at bases and tips of filopodia during formation (*t_0_*) and throughout their lifespan (*t_n_*). In addition, fluorescence at the boundary is recorded at the position closest to the site of nascent filopodium formation for a defined number of frames prior to initiation (*t_-n_*). **(B)** Example filopodium from a growth cone (shown in panel D) expressing mNeonGreen-ENA and GAP-RFP, illustrating the accumulation of ENA signal at the advancing lamellipodial edge before and during filopodium formation (orange outlined region in B). **(C)** Quantification of mNeonGreen-ENA fluorescence in the (predicted) base area (orange) and in the tip (green) of the single example filopodium from panel B. Base fluorescence and thresholded tip fluorescence are normalised against growth cone body fluorescence. After the moment of formation (*t_0_*) the enhanced fluorescence signal travels onwards with the moving filopodium tip while dropping at the base. **(D)** A still image from a timelapse (Video 5) of a growth cone showing three filopodia during formation all exhibiting enriched levels of ENA fluorescence signal (arrowheads). **(E)** Normalized base fluorescence (mean and 95% confidence interval) at newly forming filopodia (n = 53, from 8 growth cones) expressing mNeonGreen-ENA (cyan), showing accumulation at the site of future filopodium initiation in the timepoints immediately preceding its formation; this accumulation is not observed for the membrane marker GAP-RFP (red). **(F)** A still image from a timelapse (Video 6) of a growth cone expressing mNeonGreen-VASP during the moment of filopodium formation (inset). **(G)** Normalised base fluorescence at newly forming filopodia in growth cones expressing mNeonGreen-VASP (n = 31, from 7 growth cones), showing an increase in fluorescence before initiation. **(H)** Summary of filopodia properties in neurons expressing mNeonGreen-ENA, compared to mNeonGreen control. **(I)** Summary of filopodia properties in neurons expressing mNeonGreen-VASP, compared to mNeonGreen control. See Table S1 for full information. Scale bars: 5 μm (main panels), 1 µm (inset).

Given the known functional importance of ENA/VASP proteins to filopodia formation in the *Xenopus* RGCs we asked when in the life cycle of filopodia protrusion they are recruited to growth cone filopodia. VASP has previously been reported to localize to the leading edge before filopodium formation (Disanza et al., 2013; Svitkina et al., 2003). We analysed the pre-initiation base fluorescence in a timelapse series of growth cones expressing mNeonGreen-ENA or mNeonGreen-VASP. The expression of mNeonGreen-ENA had no significant effect on length, straightness, median rates of tip extension and retraction, median rates of base invasion and retraction, the proportion of time tip spent extending, retracting or stalling, or tip persistence (Table S1 and Fig. 4 H; n = 59 [control] and 99 [ENA] filopodia from N = 7 growth cones in each condition). mNeonGreen-VASP expressing filopodia showed a 14% reduction in the median rate of tip extension (Table S1 and Fig. 4 I; n = 55 [control] and 90 [VASP] from N = 8 [control] and N = 11 [VASP] growth cones; Mann-Whitney Holm-adjusted P = 0.044), but displayed no significant effect on other measured parameters. Thus, the filopodia in mNeonGreen-ENA and VASP-expressing growth cones were largely similar to filopodia in the control, mNeonGreen-expressing growth cones.

We observed that mNeonGreen-ENA localizes to the leading edge of advancing lamellipodia immediately preceding filopodia formation (Fig. 4 B-C, Video 5). In newly detected protrusions (n = 52) from 8 growth cones (an example in Fig. 4 D shows three newly formed or forming filopodia), we observed on average a slow and modest gradual increase in membrane-proximal fluorescence at sites of future filopodia initiation (Fig. 4 E). Base fluorescence of ENA peaked at 2 s before formation, followed by a decrease upon filopodia formation as the tips move away from the base region. Such increase before formation was not observed for a membrane-localising control construct (GAP-RFP), indicating that the observed peak was not a consequence of non-specific membrane localization. A similar pattern of accumulation prior to filopodia formation was observed for mNeonGreen-VASP (Fig. 4 F-G, Videos 6-7) (Svitkina et al., 2003).

There is variability between filopodia in the extent to which such increase in fluorescence signal is observed prior to initiation. We wondered whether the extent of ENA or VASP accumulation before formation affects the subsequent properties of filopodia. To address this question, we looked at the correlation between ENA or VASP intensity before formation and each of the measured properties for all filopodia in our dataset. (The accuracy of predicted base positioning diminishes with time further away from the moment of formation; for increased accuracy, we only used the last 6 seconds before formation, which also coincides with the time where ENA and VASP accumulation was greatest on average.) Significant negative correlation was found with the proportion of time spent stalling – filopodia with stronger ENA fluorescence before formation spent less time stalling (Fig. 4 J). It is possible that filopodia which start with a surge of ENA end up with higher level of ENA overall therefore giving rise to more movement. However the correlation between ENA intensity at the tip and time spent stalling has a lower overall correlation than base intensity with time spent stalling (Fig. 4 K). Thus we find that that those filopodia that start their lives with a surge of ENA accumulation spend more time in a moving state compared to those with less ENA accumulation at the membrane immediately prior to formation, without differing in other parameters. No significant correlations between VASP accumulation before filopodia formation and parameters of filopodia dynamics were detected (n = 31, α = 0.05, with Holm-adjustment for multiple comparisons).

### Measuring protein accumulation at the tip during filopodia extension

In nascent filopodia, actin polymerization at the distal end propels the extension of the filopodia tips, driven by actin-regulatory factors that localize to the tip (referred to as the “tip complex”) (Applewhite et al., 2007; Mallavarapu and Mitchison, 1999). Our initial reconstructions of the cell boundary based on the membrane signal did not always reach all the way to the tips, meaning that the majority of tip fluorescence signal was sometimes located just outside the annotated filopodia areas (Fig. 5 A). To compensate for this we designed a refinement to tip positioning: for tip-localizing fluorescent proteins, their position can be taken into account to refine the tip coordinates positioning after they are initially assigned from the membrane marker (mapping) channel (“tip fitting”; Fig. 5 A). We could therefore monitor extensions and retractions of the filopodia tips in parallel with the fluorescence in those filopodia tips, indicative of the accumulation of proteins of interest. “Fragment joining”, an additional method for improving accuracy of filopodia reconstruction, is also included in Filopodyan: with this option, separate object fragments are extended back to the closest point on the body boundary (“fragment joining”; Fig. 5 B). Filopodia frequently move in and out of plane of focus during imaging. Those events are accompanied by sudden jumps in measured tip movement, which we filter in our R scripts for downstream processing, thus eliminating out-of-focus timepoints from analysis. In addition, in our analyses of tip fluorescence and movement, automated reconstructions were extensively manually curated to ensure highest possible accuracy.

**Figure 5.**
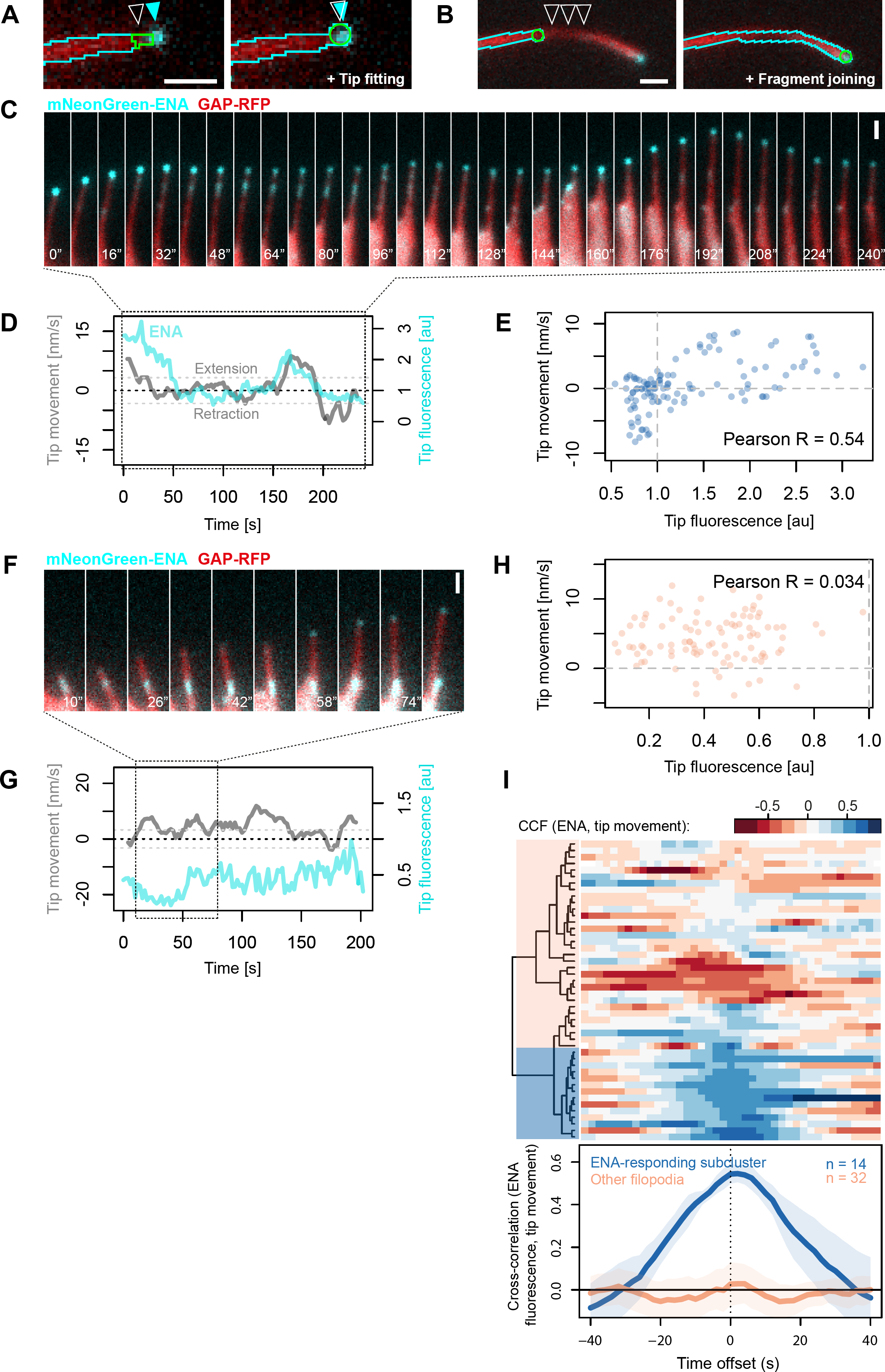
Analysis of tip fluorescence and tip movement. **(A)** For the purpose of quantifying fluorescence in filopodia tips, the ‘tip fitting’ mode of Filopodyan searches for accumulation of fluorescence signal in the immediate vicinity of initially assigned tip positions and readjusts the assigned tip position (white arrowhead) to match the position of the fluorescence signal (cyan arrowheads). **(B)** Filopodyan also allows connecting disconnected fragments, e.g. due to loss of focus (arrowheads), with the reconstructed filopodium. **(C)** A timelapse series of a filopodium from *Xenopus* RGC growth cone expressing mNeonGreen-ENA (Video 8), showing enhanced forward tip movement upon increased ENA fluorescence (at 0-20 s and 160-192 s) and absence of movement or tip retraction upon reduced ENA fluorescence (40-160 s and 192-240 s). **(D)** Concurrent tip fluorescence (normalized to growth cone fluorescence) and tip movement (smoothed with a 5-step rolling mean) measurements for the example filopodium shown in C, showing positive correlation between fluorescence and movement. Dashed grey lines at –32.5 and +32.5 nm/s represent the thresholds for retraction and extension. **(E)** Positive correlation between ENA tip fluorescence and tip movement across the timelapse for the example filopodium shown in C (with no time offset). **(F)** An example filopodium showing lack of correlation between tip ENA accumulation and tip movement (e.g. extending without ENA enrichment during 10-34 s). See also Video 9. **(G)** Concurrent tip fluorescence and tip movement measurements for the example filopodium shown in F. **(H)** Absence of correlation between ENA tip fluorescence and tip movement measurements across the timelapse for the example filopodium shown in F. **(I)** Cross-correlation (CCF) between normalized ENA tip fluorescence and tip movement for each filopodium (rows) for each value of time offset (columns) between fluorescence and movement, displayed as a heatmap. Blue: positive correlation, red: negative correlation. Negative offset: fluorescence precedes movement, positive offset: fluorescence lags behind movement. n = 46 filopodia. Blue shaded block on the dendrogram indicates the subcluster of positively correlating (“ENA-responding”) filopodia. Line plot: collective CCF values for each subcluster (ENA-responding filopodia and all other filopodia); lines and shading represent means ± 95% confidence intervals. Scale bars: 1 μm.

### Different relationships between ENA tip fluorescence and filopodia properties

In lamellipodia, the accumulation of VASP at the leading edge positively correlates with leading edge speed during extension (Rottner et al., 1999). In order to understand how ENA and VASP accumulation acutely affect the dynamic behavior of filopodia tips, we asked how tip fluorescence relates to tip movement once filopodia have protruded from the cell body. In some filopodia we observed a positive relationship between ENA tip fluorescence and tip movement: in those filopodia the loss of tip fluorescence was tightly paralleled by tip retraction, and increases in tip fluorescence paralleled their regrowth (either instantaneously or with a delay) (Fig. 5 C-E, Video 8). However, this was not the case with all filopodia, there are clear instances where positive fluorescence persists while the tip is retracting (Fig. 5 F-H, Video 9), illustrating that different filopodia within the same cell type can respond to tip accumulation of ENA in different ways.

To quantitatively assess the response of individual filopodia to ENA accumulation within their tips, we calculated for each filopodium the extent of correlation between tip fluorescence and tip movement across the entire timelapse. Allowing for the fact that the effect of one on the other may occur with a delay, we also assessed the correlation for various levels of delay between the two quantities (time offset) using a cross-correlation function (CCF) to assess this relationship as a function of time offset for each filopodium individually (Fig. 5 I). We grouped filopodia sharing a similar relationship between tip fluorescence and movement using hierarchical clustering (Fig. 5 I). 14 out of 46 filopodia (30%) displayed a positive correlation (CCF = 0.54) between ENA tip fluorescence and movement (Fig. 5 I). To verify whether this was a genuine effect rather than an artifact of the clustering algorithm, we randomized the order of tip movement measurements in a way that preserved their autocorrelation and repeated the clustering analysis on the randomized datasets. Randomization significantly reduced the cross-correlation between fluorescence and movement of the positively correlating subcluster (bootstrap P < 0.001; Fig. S3 A-E). This demonstrates that the positive correlations between ENA localization and tip movement are considerably stronger than would be expected at random in a dataset where ENA localization and tip movement are decoupled from each other.

To test this observation even more rigorously we used a Monte Carlo method. We modelled each filopodium separately based on its tip fluorescence and tip movement measurements, generating 10,000 Markov chain simulations for tip fluorescence and, independently of those, 10,000 for tip movement. We recorded pairwise correlations observed between these independently generated Markov chains, and compared the distribution of simulated correlation coefficients for each filopodium to the observed correlation coefficient in our real measured dataset for the same filopodium (Fig. S4). A single example illustrates a filopodium with positive correlation between tip fluorescence and movement (Fig. S4 A-B) and one example simulation where there is no correlation (Fig. S4 C-D). The entire set of simulations for this filopodium shows no correlation between tip movement and fluorescence, in contrast to the observed behavior (Fig. S4 E). Not a single simulation meets the correlation found in the observed data (Fig. S4 E).

The results of this analysis for each of the filopodia in the dataset showed that in 9 out of 42 filopodia, fewer than 100 in 10,000 simulations (P < 0.01) recapitulated the correlation observed in the real dataset (Fig. S4 F). (The observed correlation coefficients for these filopodia were all at least 2.8 SD away from the mean of the simulated correlation coefficients; Fig. S4 F) This means that a subset of filopodia expressing mNeonGreen-ENA displayed correlations between ENA fluorescence and tip movement at levels that would be unlikely to arise at random from their fluorescence and movement properties if these quantities were fully independent from each other, even when their temporal statistics are recreated with a Markov model.

To determine whether filopodia that have the strongest positive correlation between the accumulation of ENA in their tips and tip movement differ in other properties compared to less-correlating filopodia, we selected the top correlating cluster of filopodia for further analysis (Fig. 5 I). We call these ‘ENA-responding’ filopodia. This encompasses the possibilities that ENA accumulation influences or responds to movement, or that both are influenced by a third parameter. We compared all measured morphodynamic properties of the ENA-responding filopodia compared to all other filopodia. We found no significant differences in any of those properties between ENA-responding filopodia and other filopodia (Table S1). They also did not differ in their fluorescence properties: neither tip fluorescence nor total filopodium fluorescence or body fluorescence (Table S1), indicating that filopodia with different ENA responsiveness did not importantly differ in their level of expression of exogenous mNeonGreen-ENA.

### VASP-responding filopodia have distinct dynamic properties

Similarly for VASP, we discovered that some filopodia show a strong positive response to VASP fluorescence accumulation in their tips, whereas others do not (Fig. 6 A-F). With hierarchical clustering according to their cross-correlation between fluorescence and movement, we identified 9 VASP-responding filopodia among our dataset of 76 (12%) that were strongly responsive to VASP accumulation in their tips (CCF = 0.80) (Fig. 6 G). As for the ENA dataset above, the significance of the strength of correlation was confirmed with randomization analysis (Fig. S3 F-J) and with Markov chain simulations (Fig. S4 G). Unlike ENA-responding filopodia, which were similar to other filopodia in all measured properties, the VASP-responding filopodia reached significantly greater maximum lengths than other filopodia, had a higher median rate of tip extension, spent more time retracting and less time stalling, and their bases spent more time invading (Table S1; Fig. 6 I-J). VASP-responding filopodia and filopodia not responding to exogenous mNeonGreen-VASP coexist side-by-side within the same growth cone (Fig. 6 H, Video 10), suggesting that this observation is not due to differences in how different cells respond to exogenous VASP expression, but rather reflects heterogeneity in the molecular properties of filopodia within a single cell.

**Figure 6.**
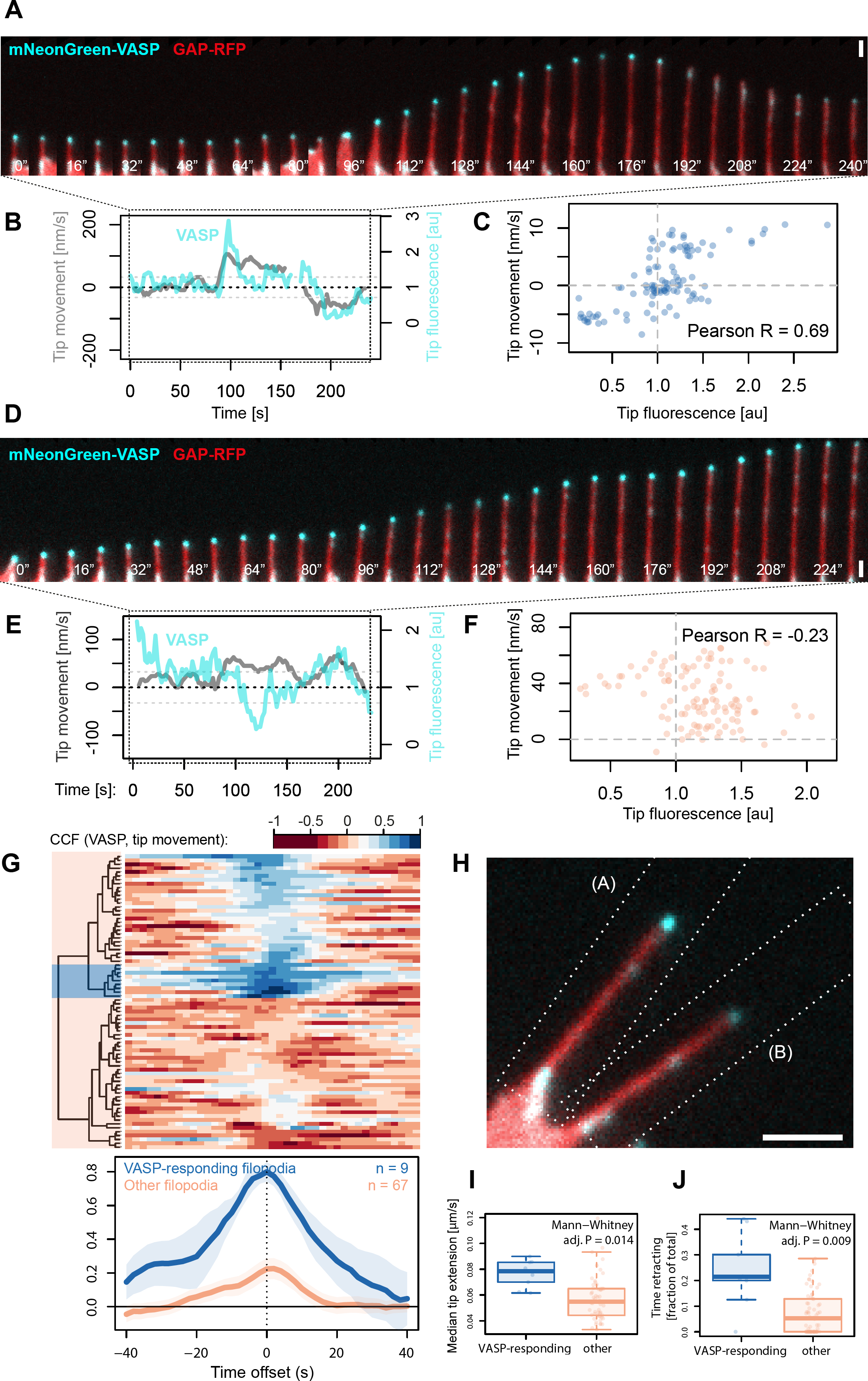
Heterogeneity of filopodia responses to VASP accumulation within their tips. **(A)** A filopodium from *Xenopus* RGC growth cone expressing mNeonGreen-VASP, showing a positive response to VASP accumulation within its tip (surge of forward movement from 96 s, retraction concomitant with the loss of tip fluorescence from 184 s). Scale: 1 μm. **(B)** Concurrent measurements of VASP fluorescence and tip movement for the example filopodium shown in A. (Interruption at t = 160 s represents loss of focus during those timepoints.) **(C)** Positive correlation between VASP tip fluorescence and tip movement across the timelapse for the example filopodium shown in A (at time offset 0). **(D)** An example filopodium expressing mNeonGreen-VASP showing stalling at elevated levels of VASP (0-80 s) and forward tip extension during a period of reduced VASP fluorescence (112-136 s). **(E)** Concurrent VASP fluorescence and tip movement measurements across the timelapse for the filopodium shown inD. **(F)** Absence of positive correlation between VASP fluorescence and tip movement across the timelapse for the filopodium shown in D, at time offset 0. **(G)** Cross-correlation between VASP fluorescence and tip movement for each filopodium (rows) as a function of time offset (columns). n = 76 filopodia. Red: negative correlation, blue: positive correlation. Negative offset: fluorescence leads before movement, positive offset: fluorescence lags behind movement. Blue shaded block over the dendrogram represents most strongly positively correlating (“VASP-responding”) filopodia, red indicates all other filopodia. **(H)** The VASP-responding and other filopodia from panels A and D occupy adjacent positions in the same growth cone. Dashed rectangles indicate crop regions used for the kymographs in panels A and D. (A still image from Video 10.) ( **I-J)** Filopodia from the VASP-responsive subcluster have a higher rate of tip extension (I) and spend more time retracting (J) than other filopodia. P values are provided for Mann-Whitney test after Holm correction for multiple comparisons. Full information on subcluster properties is in Table S1. Scale bars: 1 μm (A, D), 2 μm (H).

In our analysis subgroup allocations according to VASP responses depend on manual selection of clustering thresholds. To assess the robustness of our observations, we repeated the analysis so that a larger group of filopodia was assigned to the VASP-responding cluster (n = 37 out of 76 filopodia, 49%; Fig. S5 A). In this case, their collective CCF was reduced compared to the smaller subset of most strongly positively correlating filopodia, as expected (CCF = 0.6 compared to CCF = 0.8 for the smaller subset of 9 filopodia), but still peaked at offset = 0 (Fig. S5 B). Importantly, the correlation was strongly significant according to the randomization analysis - 0 out of 1000 randomizations recapitulated the observed cross-correlation given the defined cluster size (Fig. S5 C). We repeated the morphological and dynamic assessment of whether this expanded subgroup of filopodia had the same distinct properties as the smaller subgroup. While the maximum length and mean tip extension rate were no longer significantly different between the larger VASP-responding subgroup and other filopodia, the increase in time spent retracting and the decrease in stalling time compared with the other filopodia were preserved (Table S1).

## Discussion

### Automated detection and analysis of filopodia

We developed Filopodyan, an open-source image analysis pipeline for automated segmentation and analysis of filopodia, which rapidly annotates and tracks large numbers of dynamic filopodia. It provides options for interactive manual correction of automated reconstructions, allowing its use even in situations when accurate segmention of filopodia is difficult. We demonstrate that Filopodyan robustly identifies filopodia in a variety of model systems: two neuronal cell types in culture (RGC growth cones and dendritic filopodia in human cortical neurons) and two different cell types during *Drosophila* development *in vivo*, and across a range of imaging modes (TIRF/HILO, two-photon, and conventional line-scanning confocal microscopy) (Fig. 2). Filopodyan has therefore proven useful in various contexts important for the study of filopodia, and its flexible format means it is widely adaptable. Filopodyan is written in Fiji and R and all plugin and analysis scripts are publicly available on https://github.com/gurdon-institute/Filopodyan.

In addition to measuring filopodia shape and movement, our plugin measures the fluorescence of the whole filopodium, the tip and base, as well as the predicted base position before filopodium formation. This enables the analysis of the roles of specific proteins on the scale of seconds and micrometers, which is necessary to understand the molecular basis of filopodium growth and behavior at the relevant spatiotemporal scale of protrusive behaviour.

### Advantages and disadvantages of Filopodyan

Filopodyan measures filopodia fluorescence with full automation and generation of systematic large datasets of fluorescence intensity at membrane-proximal sites during formation and at the tip. Segmentation uses a membrane marker, providing an alternative to the actin signal used by FiloQuant (Jacquemet et al., 2017, *Preprint*) and CellGeo (Tsygankov et al., 2014). Filopodyan has the flexibility to allow high-throughput (batch) processing or a highly user-interactive mode that enables generation of manually curated datasets. Tip tracking and fragment joining for enhanced accuracy of measurement at filopodia tips are significant features compared to previous software. Filopodyan has wider scope than FiloDetect (Nilufar et al., 2013), and is implemented in ImageJ rather than commercial software, MATLAB. Filopodyan is more tailored for filopodia specifically compared to MATLAB-based CellGeo (Tsygankov et al., 2014) and ImageJ-based ADAPT (Barry et al., 2015), where filopodia are a subset of protrusions that are capable of being detected. FiloQuant is best suited for filopodia within 3D environments, and does not incorporate protein fluorescence measurements (Jacquemet et al., 2017, *Preprint*). Galic and colleagues have pioneered systematic fluorescence measurement within filopodia, looking at both tip fluorescence and fluorescence along the length of the filopodium, in low-throughput (Saha et al., 2016). Filopodyan measures the fluorescence at the membrane-localized sites of actin incorporation at both tips and bases, and links this with the parameters of filopodia morphology and dynamics in large datasets, in open source format and with a user-friendly GUI.

Filopodyan performs well on high-magnification, high-quality images in cell types with clearly separated filopodia, in two-dimensional time series. The way in which Filopodyan assigns the tip (furthest point away from the base at the cell body) means that branched filopodia tip positions are inaccurately assigned. Here we have manually removed such filopodia from further analysis and recommend that Filopodyan is not used in automated mode when branched filopodia are prominent. The segmentation process used by Filopodyan is based on erosion-dilation (similar to previous software ADAPT and FiloDetect) and is not appropriate for overlapping or looping filopodia. Different segmentation strategies, for example those used by FiloQuant (Jacquemet et al., 2017, *Preprint*) may perform better in these cases. While filtering steps and manual editing help with cells that have overlapping, branching or looping filopodia, Filopodyan has not been developed in such situations.

### Mechanisms of filopodia initation

A longstanding debate in filopodia formation is whether filopodia arise from and are embedded within a dynamic actin network of the lamellipodium (‘convergent elongation’) or whether they form anew independently of the preexisting network (‘tip nucleation’) (Yang and Svitkina, 2011). It is clear that cells are capable of producing filopodia through diverse mechanisms – for instance, in fibroblasts lacking ENA/VASP proteins or formins mDia1/mDia2, expression of either VASP or mDia2 results in overproduction of filopodia, but they have different dynamic and structural properties (Barzik et al., 2014). Similarly during *Drosophila* dorsal closure, Enabled and Diaphanous drive the production of filopodia with distinct properties *in vivo* (Nowotarski et al., 2014). To understand how cells normally produce filopodia and to assess the involvement of various actin regulators during their formation, it is necessary to systematically quantify their accumulation at the membrane. This has so far only been reported in a cell-free system of filopodia-like structures on supported lipid bilayers, suggesting a hierarchical order of protein accumulation to newly forming filopodia (Lee et al., 2010). We used Filopodyan to quanitify the enrichment of ENA and VASP at the membrane prior to filopodia initiation and protrusion in growth cones of RGC axons (Fig. 4), as a first step towards characterization of the relative timing of recruitment of actin-regulatory and membrane-binding proteins to filopodia initiation sites. While *Xenopus* RGCs offer a powerful context for studying filopodia of characterized function, one limitation is the need for exogenous expression of labeled proteins. Importantly, the expression of mNeonGreen-ENA or VASP did not substantially alter the morphodynamic properties of filopodia (Fig. 4 H-I; Table S1), indicating that the level of exogenous protein expression was not disruptive to filopodia at the levels we have used. Emergent gene editing technology is offering a way to label proteins endogenously and further refine such quantitative approaches.

### Molecular mechanisms of filopodia tip extension

We developed a method to quantify the relationship between tip fluorescence and filopodia protrusion by cross-correlation and clustering analysis, similar to previous approaches employed for lamellipodia (Barry et al., 2015; Lee et al., 2015; Machacek et al., 2009). We show that the accumulation of ENA and VASP within filopodia tips can positively correlate with their extension (Fig. 5 and 6). This effect is limited to a subset of filopodia, whereas the extension of other filopodia appears independent of the level of ENA or VASP fluorescence in their tips. While we cannot exclude a potential contribution of endogenous unlabeled ENA or VASP, growth could also be under the control of other actin regulatory proteins, such as formins. While ENA-responding filopodia harbour similar properties to other filopodia, the most positively VASP-responding filopodia grow to greater lengths, exhibit faster tip extension and spend more time in a dynamic state than other filopodia. The larger subgroup of filopodia showing VASP-responsiveness (approximately half), also spend more time retracting and less time stalling than the remaining filopodia. It may be the filopodia with lower correlation between VASP and tip movement are responding to a different subset of actin regulators. To test whether the combination of actin regulators at the tip is a more relevant measure than any single one, labelling of multiple proteins at the same time and combining with molecular interventions will be needed.

In our cross-correlation analysis of tip fluorescence and tip movement, we worked under the assumption that filopodia occupy a single state of responsiveness throughout their lifetime. However, in time series of individual filopodia, we have observed cases where filopodia not responding to VASP fluorescence seemed to switch to a responding state (e.g. Fig. 6 E, after t = 170 s). Longer time periods of data acquisition are needed to determine whether such switching is a common feature of filopodia.

### Filopodia in brain development

During their navigation *in vivo*, axons respond to a number of extracellular signals. Many of these signals have been shown to affect filopodia formation, including glutamate (Zheng et al., 1996) and the classical guidance cues Netrin-1 (Lebrand et al., 2004), and Slit (McConnell et al., 2016). Differences in the molecular mechanisms and morphological and dynamic parameters are likely tuned by the complex signaling environments in the brain. This in turn could dictate microtubule capture and stabilization of filopodia that support neurite formation (Geraldo et al., 2008), growth cone turning (e.g. (McConnell et al., 2016) or the formation of new branches in search of synaptic partners (Kalil and Dent, 2014). Filopodyan thus provides a tool to link molecular regulation of filopodia to the cellular dynamics of brain development.

## Acknowledgements

Authors would like to thank the Gallop and Harris/Holt labs for insightful suggestions, Ulrich Dobramysl, David Jordan and Edouard Hanezzo for discussions on data analysis, and Edouard Hanezzo and Ulrich Dobramysl for critical reading of the manuscript. Jennifer L Gallop and Vasja Urbančič are supported by the Wellcome Trust (WT095829AIA), Julia Mason and Benjamin Richier by the European Research Council (281971), Christine Holt by the Wellcome Trust (Programme Grant 085314) and the European Research Council (Advanced Grant 322817). The Gurdon Institute is funded by the Wellcome Trust (203144) and Cancer Research UK (C6946/A24843). The authors declare no competing financial interests.

## Author contributions

V. Urbančič, C.E. Holt and J.L. Gallop conceived the study and designed the experiments. J. Mason created the DNA constructs. V. Urbančič conducted experiments (Figs. 2A and Figs. 3-6) together with B. Richier (Fig. 2B-C) and M. Peter/F.J. Livesey (Fig. 2D). R. Butler developed the Fiji code. V. Urbančič developed the R code and data analysis. V. Urbančič and J.L. Gallop wrote the manuscript with contributions from all authors.

## Methods

### Constructs & capped RNA synthesis

pCS2-mNeonGreen-FA was generated by amplifying mNeonGreen (Shaner et al., 2013) from pNCS-mNeonGreen (supplied by Allele Biotechnology & Pharmaceuticals) with primers containing EcoRI and FseI sites (fwd: GCATGAATTCACCATGGTGAGCAAGG, rev: GATCGGCCGGCCTCTTGTACAGCTCGTCC), subsequently cloning the PCR product into pCS2-His-FA; pCS2-His-FA was constructed by digesting pCS2-FA with FseI and AscI and annealing 5’-phosphorylated linker oligos containing His_6_, EcoRI and FseI sites (fwd: TTACCATGCATCATCATCATCATCACGAATTCAGGCCGGCCTGAGG, rev: CGCGCCTCAGGCCGGCCTGAATTCGTGATGATGATGATGATGCATGGTAA CCGG). mNeonGreen-ENA was generated by amplifying the *Xenopus laevis* ENA sequence (BC073107) from pCMV-Sport 6-ENA (Open Biosystems) using primers containing FseI and AscI sites (fwd: GCATGGCCGGCCACCATGAGTGAACAGAGCATC, rev: GGCGCGCCCTATGCGCTGTTTG), and cloning into pCS2-mNeonGreen-FA using FseI and AscI. mNeonGreen-VASP was generated by amplifying the *Xenopus laevis* VASP sequence (BC077932.1) from His-mCherry-VASP (Lee et al., 2010) with primers containing containing FseI and AscI sites (fwd: GCATGGCCGGCCACCATGAGTGAGACAGC, rev: GGCGCGCCGGTCAAGGAGTACCC), subsequently cloned into pCS2-mNeonGreen-FA using FseI and AscI. GAP-RFP (Lin et al., 2009) was a gift from Holt laboratory. For capped RNA synthesis these plasmids were linearised with NotI, and transcribed *in vitro* using SP6 mMessage mMachine (Ambion) following manufacturers instructions, and diluted in RNase-free water.

### *Xenopus* retinal ganglion cell growth cones

*Xenopus* embryos were obtained by *in vitro* fertilisation and reared in 0.1 x modified barth’s saline (MBS) at temperatures ranging from 14 to 18 ^o^C. Developmental stages were determined according to Nieuwkoop and Faber (Nieuwkoop and Faber, 1967). At embryonic stages 26-28, RNA was introduced into eye primordia by eye electroporation at 0.5 μg/μl per construct as previously described (Falk et al., 2007). Briefly, embryos were placed inside custom-made Sylgard electroporation chambers with platinum electrodes placed either side of the embryo. RNA was delivered by microinjection to a site medial to eye primordia, followed by the application of eight 18mV pulses of 50 ms duration, separated by 1 s intervals. Eye explants of stage 35-36 embryos were cultured as previously described (Leung and Holt, 2008), on 35mm glass-bottom culture dishes (Ibidi) coated with 10 μg/ml PLL overnight (Sigma) and with 10 μg/ml laminin (Sigma) for 3-5 hours(Leung and Holt, 2008). Eye explants were cultured in phenol red-free L-15 culture media (Invitrogen), and imaging was conducted 19-23 hours after plating. Imaging of retinal explants was conducted in the same culture media with highly inclined laminated optical sheet (HILO) illumination (Tokunaga et al., 2008) on a custom-made TIRF setup based on a Nikon Eclipse Ti-E inverted microscope equipped with an iLas2 illuminator (Roper Scientific), a CMOS camera (Hamamatsu Flash 4) and an Optosplit beam splitter (Cairn Research). Images were acquired in a single z plane at a rate of 2 s per timepoint with a 100x 1.49 NA oil immersion objective at room temperature, using MetaMorph software (Molecular Devices). Only growth cones clearly separated from other axons were chosen for imaging.

### *Drosophila* stocks and embryo live imaging

Flies were raised at room temperature on standard fly food. The following fly stocks were obtained from the Bloomington Drosophila stock center: {en2.4-Gal4}e16E (BL30564); w; P{UAS-mCD8.ChRFP} (BL 27392). w; btl-Gal4 UAS-CAAX-mCherry/TM6b was obtained from Shigeo Hayashi. Genotypes used for live imaging were *;en-Gal4/CyO; UAS-cd8mCherry* for leading edge cells during dorsal closure and *btl-Gal4 UAS-Cherry-CAAX/TM6b* for terminal tracheal cells. After egg-laying overnight at 25°C, embryos were dechorionated in thin bleach for one minute and mounted on a coverslip with heptane glue and covered with water. Embryos at the end of stage 14 and at stage 16 were selected for imaging dorsal closure and tracheal cells, respectively. Live imaging was performed at room temperature on an inverted Leica TCS-sp5 confocal microscope equipped with a 63x 1.4NA plan Apo oil immersion objective and Hybrid detectors HyD, using Leica software. Stacks of 7-10 z sections (0.5 or 0.7µm Z step size) were taken every 15 seconds for 10 minutes. Using ImageJ, a maximum projection was then applied to use the plugin Filopodyan for filopodia reconstruction.

### Human iPS cell lines, culture and imaging

Cortical differentiation: All cells were maintained at 5% CO2 at 37°C in a humidified incubator. NDC1.2 (Israel et al., 2012) iPSC were grown in Essential-8 medium (ThermoFisher) as feeder free cultures on Geltrex (ThermoFisher) coated plates. Neuronal induction was performed as described previously (Shi et al., 2012). In brief, iPSCs were passaged with 0.5 mM EDTA and plated at high density to reach 100% confluency in 24 h when neuronal induction was started (day 0). Essential-8 medium was changed to neuronal induction medium consisting of neuronal maintenance medium supplemented with 10μm SB43152 (Tocris) and 1μm Dorsomorphin (Tocris). The medium was changed daily until reaching day 12. On day 12 the neuroepithelial sheet was lifted off with Dispase (ThermoFisher), broken up to smaller clumps and plated on laminin (Sigma) coated plates in neuronal induction medium. On day 13 the medium was changed to neuronal maintenance medium supplemented with 20 ng/ml FGF2 (PeproTech). Medium was changed every other day and FGF2 was withdrawn from the medium on day 17. Cells were split with Dispase at a 1:2 ratio when neuronal rosettes started to meet. On day 25 cells were disassociated with Accutase (ThermoFisher) and re-plated on laminin coated plates.

Until day 34 cells were expanded at a 1:2 ratio when they reached 90% confluency. On day 36 neurons were plated on laminin coated plates at 85000 cells/cm^2^ and used for subsequent experiments. Neuronal maintenance medium (NMM) (1L) consists of 500 ml DMEM:F12 + glutamax (ThermoFisher), 0.25 ml Insulin (10 mg/ml, Sigma), 1 ml 2-mercaptoethanol (50 mM ThermoFisher), 5 ml non-essential amino acids (100x ThermoFisher), 5 ml Sodium Pyruvate (100 mM, Sigma), 2.5ml Pen/Strep (10000 U/μl, ThermoFisher), 5 ml N2 (ThermoFisher), 10 ml B27 (ThermoFisher), 5 ml Glutamax (100x, ThermoFisher) and 500 ml Neurobasal (ThermoFisher) medium. Two-photon imaging of human neuronal filopodia: Human cortical neurons were transfected on day 38 with a plasmid expressing NeonGreen fluorescence protein from the human synapsin 1 promoter using Lipofectamin3000 (ThermoFisher) following the manufacturers protocol. Neurons were kept in neuronal maintenance medium until they reached day 88. To reduce background fluorescence during two-photon imaging the medium was changed to neuronal maintenance medium without phenol red DMEM:F12 + L-glutamin – phenol red (ThermoFisher). The inverted two-photon microscope (VIVO multiphoton, 3i) was equipped with a humidified incubator and we performed imaging at 5% CO2 at 37°C. NeonGreen was excited at 950 nm using a tunable pulsed laser (InSight DS+, Spectra Physics) and timelapse image stacks of filopodia were taken over 30 min at a 2 min time interval using a 63x objective (X/Y pixel size: 0.053 μm, Z step size: 0.38 μm). Image stacks were exported to tiffs using Slidebook (3i) software.

### Segmentation and tracking in ImageJ/Fiji

Full code of our plugin is publicly available at https://github.com/gurdoninstitute/Filopodyan. Images are first processed with a Laplacian of Gaussian filter to reduce noise and enhance contrast of the features of interest.

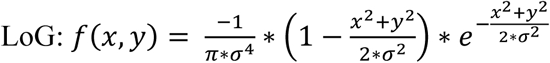

The resulting images are binarised using a choice of auto-thresholding algorithms, and the largest object in the field of view is assumed to be the growth cone/cell body. The binary mask at each time point is segmented into the growth cone/cell body and filopodia by applying n erode operations, where n is the number required to remove all filopodia from the mask, followed by n dilate operations to restore the size of the resulting growth cone body mask. This final mask is subtracted from the original mask to leave only separate filopodia for analysis. For each of the segmented filopodia, ROIs are calculated at the base and tip defined by intersection with the growth cone body and the point furthest from it respectively. Filopodia are tracked over time using a rapid one-step Hungarian linear assignment algorithm, with linking costs calculated using the following equation:

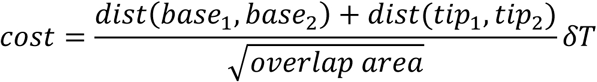

Tracks can be filtered using several different threshold values, and tracking errors in the resulting reconstruction can be manually corrected using a simple interface to delete filopodia from tracks and draw the desired links.

### Measurement of metrics describing filopodial shape, dynamics and fluorescence

*Length*– An estimate of length is defined in Filopodyan coarsely as one half of perimeter of the protrusion ROI. Because this leads to an overestimation of length for short filopodia, we adjusted our perimeter length computation with a correction that substantially reduces the error in length estimation for short filopodia. This correction accounts for the contribution of base width and tip roundedness to perimeter length, and subtracts an estimate of these quantities multiplied with a scaling factor. Thus, estimated length is defined as:

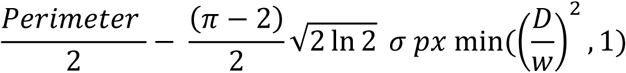

This is derived from protrusion ends-correction (tip correction: ¼ circumference with radius *w*/2, and base correction *w*/2); width *w* is estimated for this purpose by the formula for full width at half maximum using user-defined σ of Laplacian of Gaussian (provided during segmentation step) and pixel width 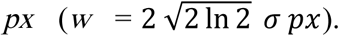 Scaling factor min((*D/w*)^2^, 1) adjusts the degree of correction for protrusions whose width is greater than their Euclidean (base, tip) distance (*D*).

*Straightness*– Defined as:

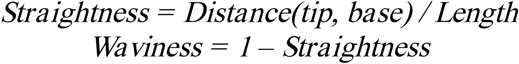

Straightness is underestimated for short filopodia (and equally, waviness is overestimated for short filopodia), in practice not reliable for short structures. The plugin is not intended to look at branched structures. If applied to branched structures, straightness and waviness will be inaccurate as a measure of actual projection straightness and will instead reflect the degree of branching.

*dL* **–** change in length between successive timepoints. Defined as:

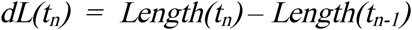

dL is affected by tip movement as well as base movement, which are more informative metrics, hence dL was largely ignored in our analysis. dL could be useful when a measure of length change independent of corrections for position or direction is required.

*Tip movement*– rate of directional tip extension or retraction, providing a direct measure of the rate of tip extension or retraction between successive timepoints. Defined as direction-corrected tip movement (‘DCTM’) = tip displacement from the preceding timepoint, projected onto the vector connecting base and tip at current timepoint:

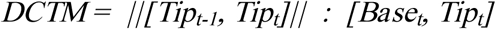

DCTM is closely related to tip speed and equals tip speed when the angle of filopodium is constant between successive timepoints. When the angle of a filopodium changes between successive timepoints (e.g. due to sideways movement of the tip), DCTM is corrected for that change in angle (e.g. so as not to overestimate extension if increased speed was due to swinging of the filopodium). DCTM equals dL when the base is static and filopodium angle is constant. DCTM can be conceptualized as tip speed within a simplified one-dimensional representation of the filopodium. We believe this approach is the best available readout for the rate of productive polymerisation of actin filaments at filopodia tips during extension, or their breakdown during retraction, assuming no retrograde flow.

Filopodyan outputs a wide range of metrics representing the behavior of filopodia, and we anticipate that each user will wish to select those of interest for their own experiments and apply their own choice of statistical analysis. We applied downstream analysis to the output tables using scripts in R for our pipeline, but any program could be used depending on the required functions. These scripts apply a tip movement filter to reject measurements outside 0.5-99.5 percentile range (often caused by out of focus effects and reconstruction errors), smoothed with a 5-step rolling mean, and further divided into:

*Tip extension rate* = DCTM where DCTM > 32.5 nm / s

*Tip retraction rate* = DCTM where DCTM < -32.5 nm / s

*Initial tip movement* = median DCTM over first 10 timepoints (0-20 s)

*Base movement* **–** rate of directional base invasion or retraction, defined as:

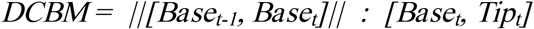

Direction-corrected base movement, directly analogous to direction-corrected tip movement as described above. DCBM is further divided into:

*Base invasion rate* = DCBM where DCBM > 32.5 nm / s

*Base retraction rate* = DCBM where DCBM < 32.5 nm / s

*Initial base movement* = median DCBM over first 10 timepoints (0-20 s)

*Tip persistence* **–** a measure of consistency of tip movement across time

For each filopodium, the autocorrelation function (ACF) is computed for the time series of its tip movement. ‘Tip persistence’ is the root of this function (ACF of DCTM), i.e. the time required for autocorrelation of tip movement for a filopodium to drop to zero.

*Time extending / retracting / stalling* **–** proportion of time the tip spent in any of the three states of movement. Extension time: consistent extension by more than 32.5 nm/s for 10 s; retraction time: consistent retraction by more than -32.5 nm/s for 10 s. Stalling time: tip movement between -32.5 and 32.5 nm for 10 s. As in tip movement above, tip movement was smoothed with a rolling mean filter (step size = 5) prior to the calculation to reduce the effects due to measurement noise.

*Proj Mean*– Mean fluorescence intensity within the area reconstructed as protrusion (‘projection’) (cyan in Filopodyan overlay), in the channel set as measurement channel.

*Base Mean*– Mean fluorescence intensity within the base area of the protrusion (orange in Filopodyan overlay), in the channel set as measurement channel.

Predicted base area: predicted base position is defined as position on current boundary with minimum distance from the base at the preceding timepoint.

*Tip Mean* **–** Mean fluorescence intensity within the area reconstructed as tip of protrusion (green in Filopodyan overlay), in the channel set as the measurement channel. Tip is defined as the point at the boundary with maximum distance from the base. The tip may be incorrectly assigned in backwards turning, looping or buckling filopodia.

*Tip Thresholded Mean*– Mean fluorescence intensity within Otsu-thresholded tip ROI, in the channel set as measurement channel. This measure is useful when a significant proportion of the tip area has the same fluorescence intensity as signal background, separating tip signal from background.

*Body Mean*– Mean fluorescence intensity within the body area of the cell or growth cone (magenta in Filopodyan overlay), in the channel set as measurement channel.

### Additional optional functionalities in Filopodyan

#### Tip Fitting

Tip fitting to the measurement channel is done in a radius of 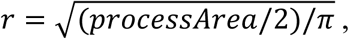, where 3 < r < 20 pixels. The tip position is set to the intensity weighted mean of the maxima coordinates and the new tip radius is set to 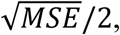 where MSE is the mean squared error of the maxima coordinates.

#### Fragment joining

Fragment joining is a useful option when parts of filopodia contain weaker signal or are partially out-of focus. Fragment joining ensures all disconnected fragments are connected to the growth cone by finding the shortest line connecting any two points on the boundaries and adding it to the binary mask. It connects fragments that are close together, then connects the joined fragments to the growth cone until they converge to a single structure over a maximum of 10 iterations.

#### Boundary Visualization

Signal intensity along the boundary of the growth cone is measured for each point by taking the mean value of a small internal area in the specified channel, and the local velocity is estimated based on the change in intensity at the boundary coordinates over time in the unprocessed image (inspired by Barry et al. 2015). The boundary signal intensity and velocity values can be used to generate colour-coded boundary images, kymographs showing change over time in 2D, and cross-correlation plots showing the relationship between signal intensity and boundary extension or retraction.

#### Adaptive Thresholding

The adaptive local thresholding function convolves the image with directional LoG kernels for the 8 principal directions, thresholds the resulting images separately using the selected method and combines them into a single mask with noise removed using a median filter.

### Quantification of morphodynamic properties of filopodia in *Xenopus* RGCs growth cones

For phenotypic characterization of filopodial properties, time series of *Xenopus* growth cones expressing GAP-RFP, coexpressed with mNeonGreen, mNeonGreen-ENA or mNeonGreen-VASP, were quantified with Filopodyan using GAP-RFP as the mapping channel with the following segmentation and filtering parameters: *Thresholding method:* Renyi entropy, *σ_LoG_* = 2.6, *erosion-dilation step (ED)* = 4, *tip fitting:* disabled; Filtering: *min frames* = 3, *min max length*= 1.8 μm, *min length change* = 0.1 μm, *max mean waviness* = 0.35. Except in the case of batch processing (Fig. 2, Fig. S1), manual editing was applied to remove reconstructions if they were false positives, out-of-focus, looping, branching, crossing or emanating from axon shaft, and obvious tracking errors were rectified. Correlation matrix visualizations (Fig. 3) were constructed with the R package ‘corrplot’ (Wei and Simko, 2016). Visual representations of morphodynamic phenotype are constructed from data normalized such that the median of the control dataset equals 0 and one interquartile range (IQR) of the control dataset equals 1. New filopodia were identified by downstream R scripts as the filopodia with min T > 1 and starting length < 2 µm to remove longer new reconstructions representing filopodia suddenly coming into focus. Initial speed of tip movement was defined as the median DCTM of the first 10 timepoints (20 s) in existence. Initial pre-formation fluorescence was defined as the mean of the last 3 timepoints before emergence. Full code is available at https://github.com/gurdon-institute/Filopodyan/tree/master/FilopodyanR (scripts ‘Correlations.R’ and ‘Correlations_DataInput.R’, and FilopodyanR Module 1, 2 and 3).

### Quantification of base fluorescence before filopodium initiation

Time series of mNeonGreen-ENA and mNeonGreen-VASP expressing growth cones were reconstructed with Filopodyan v20170201 using GAP-RFP as the mapping channel and the following segmentation and filtering parameters: *Thresholding:* Renyi entropy, *σ_LoG_* = 2.6, *ED* = 4, *number of base back frames* = 20 (other segmentation options disabled); Filtering: *min start frame*= 2, *min frames* = 3, *min max length* = 1.8 μm, *min length change* = 0.1 μm, *max mean waviness* = 0.38. Additional manual editing was applied as required to remove incorrect or unsuitable annotations (false positives, tracking errors, or filopodia arising from the axon shaft). Background measurements were subtracted from mean base fluorescence prior to normalising to background-corrected mean body fluorescence. Full code is available at https://github.com/gurdon-institute/Filopodyan/tree/master/FilopodyanR (scripts ‘Module 1.R’, ‘Module 1-2_BgCorrection.R’ and ‘BaseF.R’).

### Quantification of tip fluorescence

Time series of growth cones were reconstructed with Filopodyan using GAP-RFP as the mapping channel with the following parameters: *Thresholding:* Huang, *σ_LoG_* = 4.01, *ED* = 4, *tip fitting:* enabled, *fragment joining:* disabled. Filtering: *min frames* = 3, *min max length* = 1.8 μm, *max mean waviness* = 0.38. Extensive manual editing was applied to ensure all reconstructed structures in all timepoints represented meaningful measurements, removing reconstructed tracks that were false positives, out of focus, branching, looping, emanating from the axon shaft, or whose tip ROIs were incorrectly positioned relative to tip signal. Background-subtracted thresholded tip fluorescence was normalised to background-subtracted body fluorescence in the corresponding timepoint. Full code is available at https://github.com/gurdoninstitute/Filopodyan/tree/master/FilopodyanR (scripts ‘Module 1.R’ and ‘Module 1-2 BgCorrection.R’).

### Cross-correlation analysis and hierarchical clustering

For each filopodium, the value of cross-correlation function (CCF) between tip fluorescence (background-corrected by the subtraction of background signal near the growth cone boundary, and normalized to body fluorescence with equivalent background subtraction) and its DCTM (passed through 0.5^th^-99.5^th^ percentile filter and smoothed with 5-step rolling mean) was calculated in R using the R Stats function ccf(). Filopodia with fewer than 17 timepoint measurements were excluded from analysis because they were too short to generate meaningful randomisations (see below). CCFs of all filopodia were fed into a hierarchical clustering analysis using Euclidean distance between CCFs at offsets between -6 and +6 s as a similarity measure. Heatmap visualizations were generated using heatmap.2() function from R package ‘gplots’(Warnes et al., 2016). For randomisation controls, smoothed DCTM measurements for each filopodium were randomly reshuffled in blocks of 8 timepoints in order to preserve their autocorrelation (similar to the value of their tip persistence), and the CCF values were recalculated for each randomised time series using the same method. CCFs from the randomized time series were then analysed by hierarchical clustering as described above. To minimize bias arising from differences in cluster size, the resulting clusters were only accepted into the analysis if the number of filopodia in the top correlating subcluster was similar as in the original non-randomised dataset to within 15% (arbitrary cutoff). CCFs for all the randomised filopodia in the top correlating subclusters of each randomisation were recorded. Block randomisation, clustering and top correlating subcluster assessment were repeated until reaching 1000 accepted randomisation datasets. Full code is available at https://github.com/gurdon-institute/Filopodyan/tree/master/FilopodyanR (in FilopodyanR scripts ‘CCF.R’, ‘CCF_Randomisations.R’, and ‘CCF_subclusteranalysis.R’).

### Discrete time Markov chain simulations

For each filopodium its tip fluorescence (normalized background-corrected measurements) and tip movement (5-step rolling mean smoothed tip movement (DCTM)) were binned into 9 equal-sized intervals to assign Markov states. The R package ‘markovchain’ (Spedicato, 2016) was used to calculate transition probabilities between these states for each filopodium and to run 10,000 simulations (realisations) per filopodium transition matrix. The initial state of each filopodium was used as the starting state for all its corresponding simulations. A small fraction of filopodia produced invalid transition matrices (3 out of 45 in the ENA dataset, and 4 out 76 in the VASP dataset) and these were discarded from the Markov chain analysis. Full code is available at https://github.com/gurdoninstitute/Filopodyan/tree/master/FilopodyanR (‘FilopodyanR MarkovChains.R’).

### Online Supplemental Material

Supplemental Material associated with this manuscript is available on the journal website and includes: Figures S1-5, Videos 1-10, and Table S1 listing full phenotypic measurements and associated statistics. The complete data tables outputted by Filopodyan of growth cone filopodia parameters and fluorescence intensities that were used as the source of further analysis are also provided.

---

**Figure S1.**
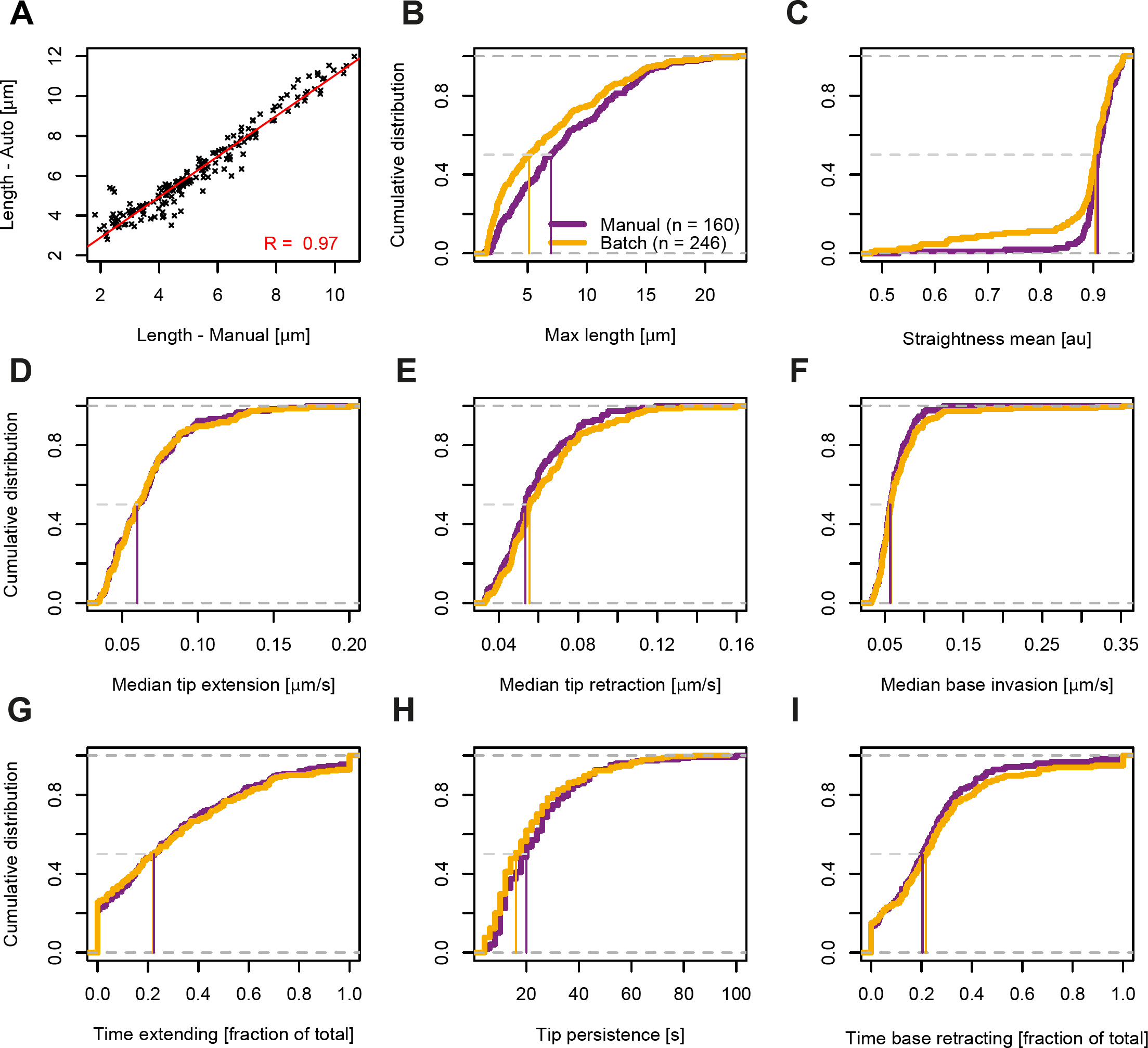
Comparison of filopodia properties measured with and without manual editing of reconstruction. **(A)** Correlation between 186 manual and automated measurements of filopodia lengths (6 filopodia, 31 timepoints each) from *Xenopus* RGC growth cone, showing a strong good positive correlation between manual and automated measurements. **(B-I)** Measurements of filopodia properties from the descriptive dataset of 19 *Xenopus* RGC growth cones. Cumulative distribution curves are shown for the filopodia dataset following automated filtering only (yellow; ‘CAD filter’) and for automated filtering followed by extensive manual curation of the reconstructions (purple, ‘Manual filter’). Median of each dataset is displayed as a vertical line intersecting the cumulative density curve at 0.5.

**Figure S2:**
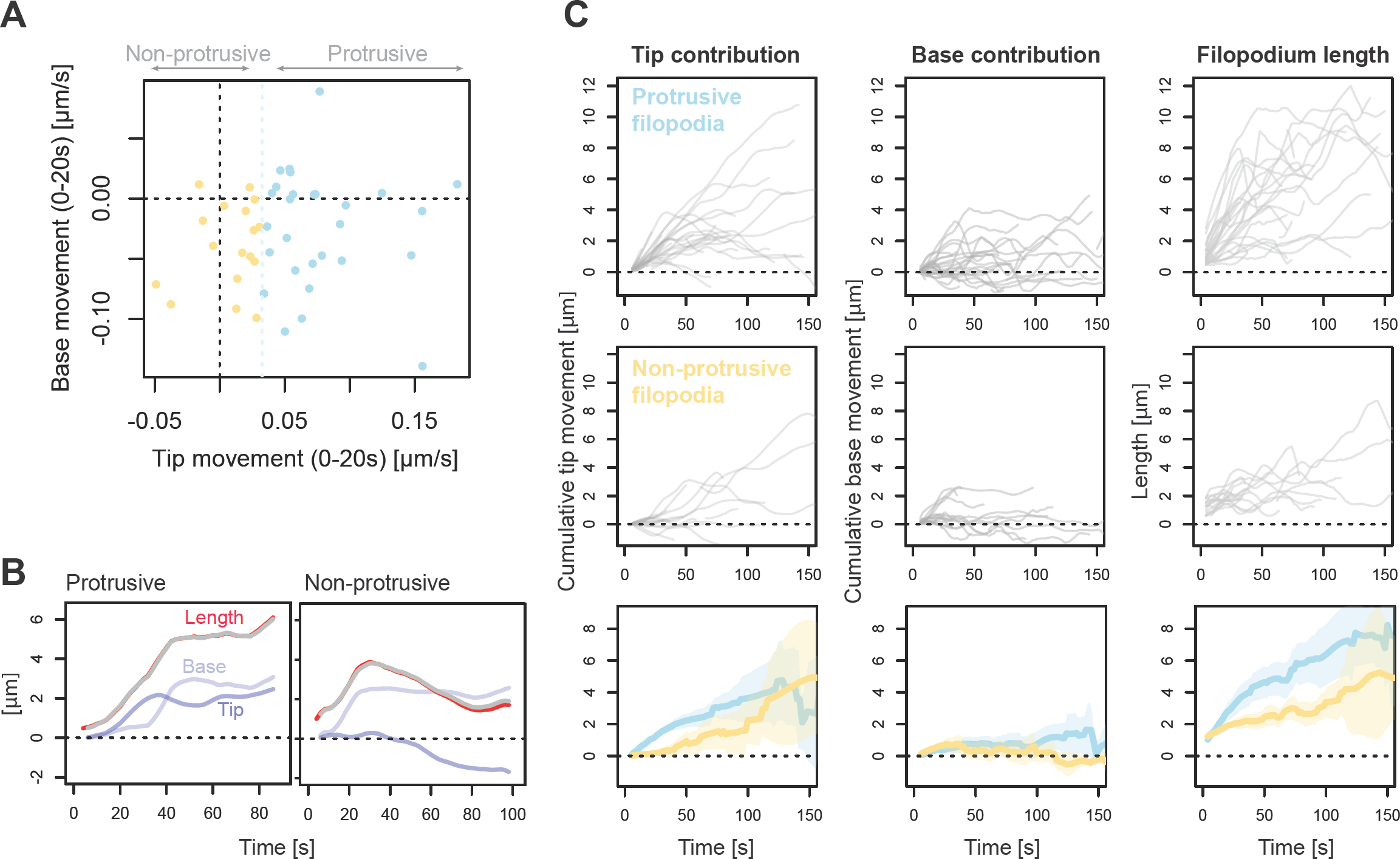
Tip and base movements in filopodia arising from protrusive or retractive events. **(A)** Scatterplot of median tip and base movement during the initial phase (first 20 s in existence) for newly forming filopodia (n = 45) in *Xenopus* RGC growth cones. Filopodia were divided into two categories depending on whether their median initial tip movement exceeded extension threshold (“protrusive”, blue, n = 28) or not (“non-protrusive”, orange, n = 17). **(B)** Examples illustrating the dynamics of tip and base movement in filopodia in each category. Red line: total filopodium length (smoothed with five-step rolling mean). Blue: contribution of tip movement to filopodium length (cumulative sum of smoothed DCTM measurements over the time series). Lighter blue: contribution of base movement to filopodium (negative cumulative sum of smoothed DCBM measurements over the time series). Grey: the sum of cumulative tip movement and cumulative base movement, adjusted for initial length at the first available timepoint, showing good overlap with filopodium length. **(C)** Line traces for cumulative tip movement (left), cumulative base movement (middle) and total length (right) over time for all newly forming filopodia in the dataset, separated according to subgroup (top: protrusive; middle: non-protrusive). Bottom row: Mean and 95% confidence interval measurements for cumulative tip movement (left), cumulative base movement (middle) and total length (right) for each of the two subgroups of filopodia (blue: “protrusive”, n = 28; orange: “non-protrusive”, n = 17).

**Figure S3.**
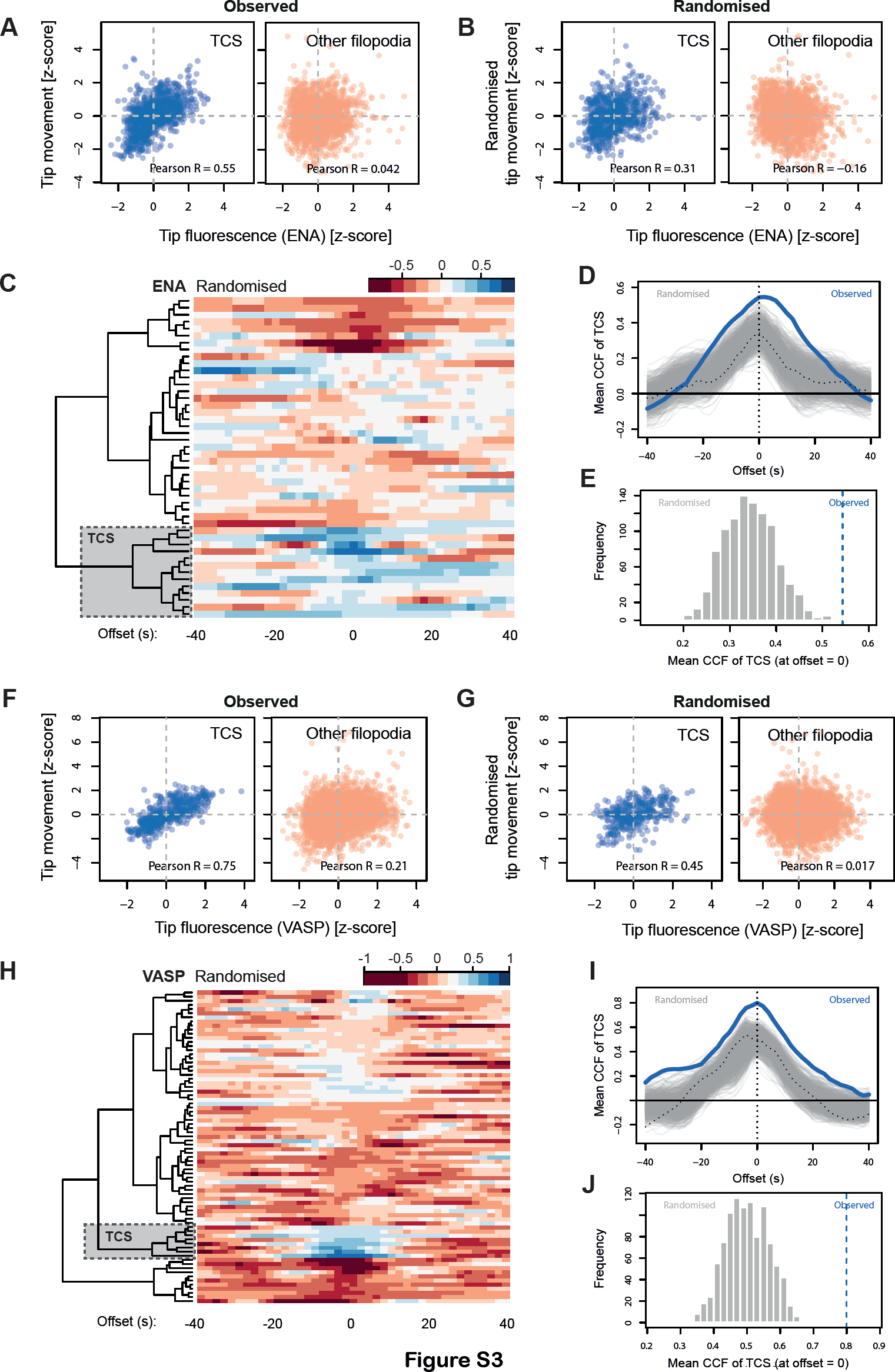
Randomisation control for the hierarchical clustering of cross-correlations between tip fluorescence and movement. **(A)** Correlation between ENA fluorescence and tip movement for ENA-responding filopodia (blue; top-correlating subcluster in the CCF heatmap in main Fig. 5 I) and for all other filopodia (red), normalized such that 0 = mean for each filopodium, and 1 = SD for each filopodium. **(B)** Correlation between ENA tip fluorescence and randomized tip movement. After hierarchical clustering on the randomized dataset (in panel c) to identify the most positively correlating subcluster (top-correlating subcluster, TCS), its correlation between fluorescence and movement is reduced by comparison to the original dataset. **(C)** Heatmap represents the level of cross correlation of tip movement and tip fluorescence for each filopodium after reshuffling the order of tip movement for each filopodium using block randomization to artificially decouple movement from fluorescence. Rows represent individual filopodia, columns indicates the offset (lag/lead) of the cross-correlation function. Grey boxed region indicates the top correlating subcluster (TCS) within the randomized dataset. Only one representative randomization out of 1,000 is shown. **(D)** Cross-correlation between (randomized) tip movement and ENA tip fluorescence in the top-correlated subcluster at different values of cross-correlation offset. Grey lines represent the top-correlated subcluster in individual randomized datasets (n = 1000), of which the representative example from panel C is indicated by dotted black line. Blue line (“Observed”) shows the cross-correlation of the top-correlated subcluster in the original, non-randomised dataset (as in Fig. 5). **(E)** Histogram showing the cross-correlation between ENA fluorescence and tip movement at zero offset for the top-correlating subcluster in all randomised datasets, with the cross-correlation of the original (“observed”) top-correlating subcluster indicated by the vertical blue line. Following randomization, none out of 1000 positively correlating subclusters are cross-correlated as strongly as the original dataset (p < 0.001). **(F-J)** As above, for VASP.

**Figure S4.**
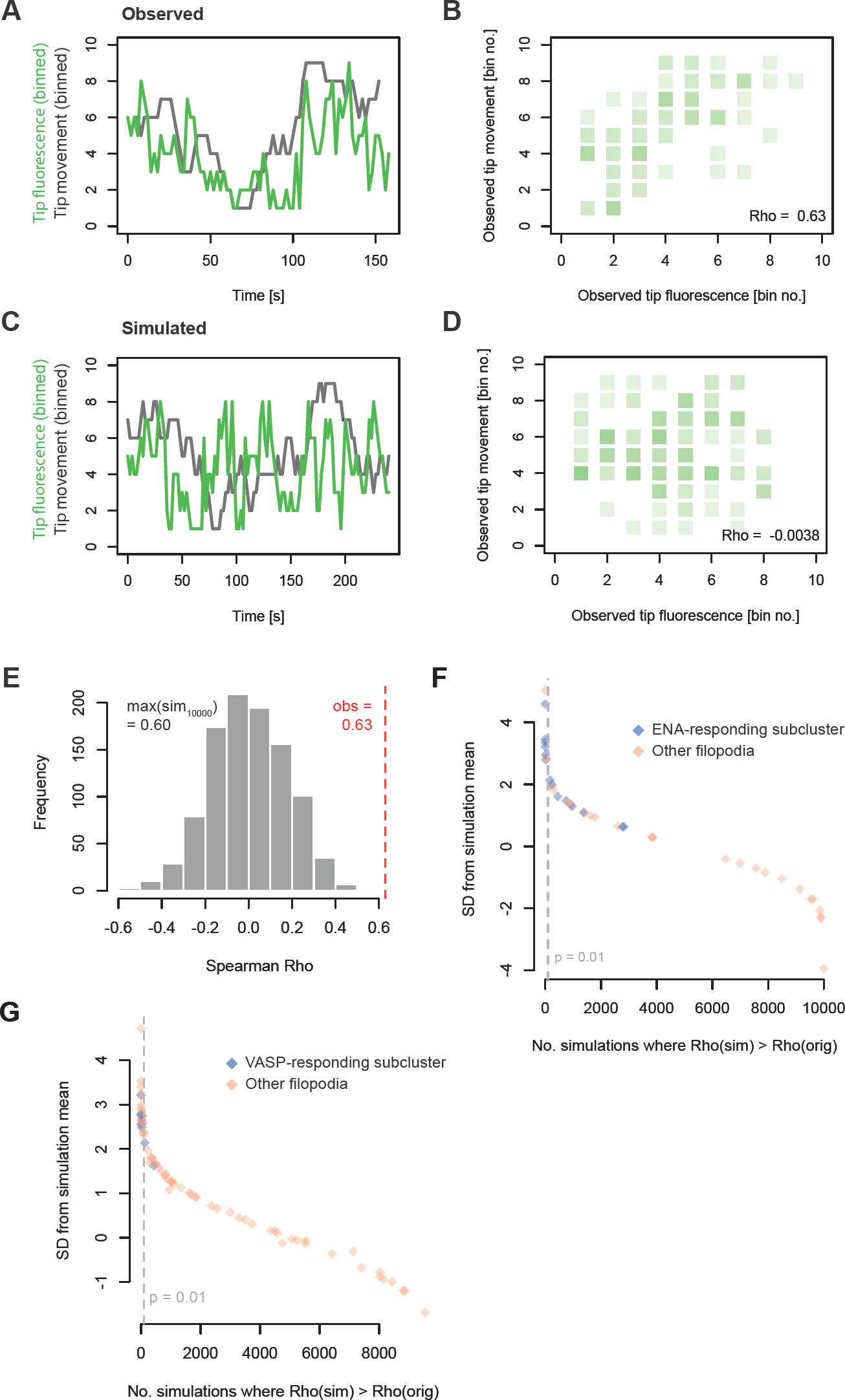
A Monte Carlo Markov chain method confirms the low likelihood of observed correlations between ENA and VASP tip fluorescence and tip movement occurring by chance. **(A)** Traces of tip fluorescence and tip movement of an example filopodium expressing mNeonGreen-ENA, binned into equal-sized intervals. **(B)** Binned fluorescence and movement measurements for the filopodium from panel A across its recorded lifetime, showing a positive correlation between tip movement and tip fluorescence for this filopodium. **(C)** A simulated filopodium dataset based on Markov chain transition probabilities derived from the real filopodium data in panels A-B. **(D)** Correlation between simulated fluorescence and simulated movement across the time series for the simulation shown in panel C. **(E)** A histogram of correlation coefficients between simulated tip movement and simulated tip fluorescence for 10,000 simulations based on the transition probabilities of one filopodium shown in A-B (including the simulation shown in panels C-D). The mean of correlation coefficients of all simulations is zero; the maximum is 0.60, lower than the correlation coefficient of the measurements in the real (observed) dataset, 0.63 (corresponding to panel B). **(F)** A summary of results of 10,000 simulations per filopodium for all filopodia in the mNeonGreen-ENA expressing dataset, providing the number of simulations whose correlation Rho (between simulated tip fluorescence and movement) exceeded the correlation Rho found in the original time series (x axis). y-axis shows the distance of the observed correlation coefficient in original time series from the mean of correlation coefficients in all simulations for that filopodium, expressed as the number of standard deviations from the mean. Colour code highlights the subcluster identity of each filopodium according to the clustering presented in Fig. 5 I. **(G)** as panel F, for mNeonGreen-VASP expressing filopodia. Colour code highlights filopodium subcluster identity according to the clustering in Fig. 6 G.

**Figure S5.**
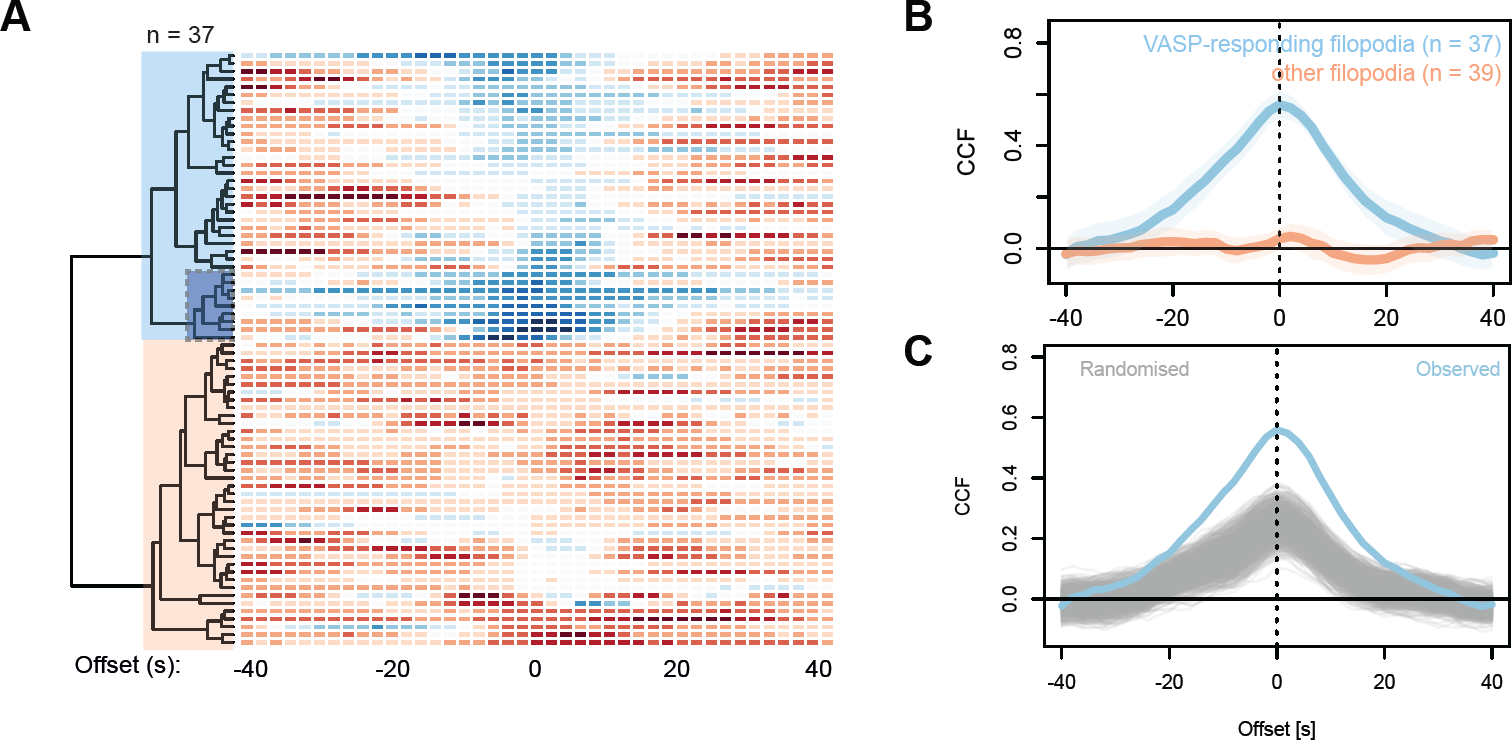
Assessment of robustness in cross-correlation against subcluster size. (A) Heatmap representing CCFs between mNeonGreen-VASP tip fluorescence and tip movement, directly duplicated from Fig. 6 G, analysed here with a different subcluster selection threshold: in the new analysis, “VASP-responding filopodia” contain a much enlarged subset of filopodia (n = 37 out of 76; light blue block over dendrogram) compared to the analysis shown in Fig. 6 G-J (n = 9 out of 76; shaded dark blue/dotted outline). (B) Mean and 95% confidence interval of the value of CCF computed collectively for VASP-responding filopodia (here, n = 37) and all other filopodia (n = 39) relative to offset. (C) Mean CCFs of the top correlating subcluster (with n size between 32 and 42) from each of 1000 randomizations of the VASP tip fluorescence and tip movement dataset. 0/1000 randomizations recapitulate the observed mean top subcluster CCF (bootstrap P < 0.001).

**Video 1.** Timelapse video of a *Xenopus* RGC growth cone, imaged with HILO illumination on a TIRF imaging system, and segmented with Filopodyan. Acquisition rate: 2 s per frame, playback speed: 12 fps. False-color look-up table (LUT) ‘Fire’: membrane marker GAP-RFP. Scale bar: 5 μm.

**Video 2.** *Drosophila* tracheal cells imaged by line-scanning confocal microscopy, segmented with Filopodyan. 15 s per frame, playback speed: 7 fps. False-color LUT ‘Fire’: membrane marker Cherry-CAAX. Scale bar: 5 μm.

**Video 3.** *Drosophila* leading edge cells during dorsal closure imaged by line-scanning confocal microscopy, segmented with Filopodyan. Acquisition rate: 15 s per frame, playback speed: 7 fps. False-color LUT ‘Fire’: membrane marker cd8-mCherry. Scale bar: 5 μm.

**Video 4.** Dendritic filopodia in induced pluripotent stem cell (iPSC)-derived human cortical neurons, imaged by 2-photon microscopy, segmented with Filopodyan. Acquisition rate: 2 min per frame, playback speed: 4 fps. False-color LUT ‘Fire’: cytoplasmic marker mNeonGreen. Scale bar: 5 μm.

**Video 5.** mNeonGreen-ENA localization during formation of new filopodia in an RGC growth cone imaged using HILO illumination. Detected filopodia were filtered for newly forming filopodia (*min start frame* = 2). Acquisition rate: 2 s per frame, playback speed: 12 fps. Red: membrane marker GAP-RFP. Cyan: mNeonGreen-ENA. Scale bar: 5 μm.

**Video 6.** Localisation of mNeonGreen-VASP before filopodia formation in an RGC growth cone imaged using HILO illumination (example from Fig. 4). Detected filopodia were filtered to newly forming filopodia (*min start frame* = 2). Acquisition rate: 2 s per frame, playback speed: 12 fps. Red: membrane marker GAP-RFP. Cyan: mNeonGreen-VASP. Scale bar: 5 μm.

**Video 7.** Localization of mNeonGreen-VASP in an RGC growth cone during formation of a single filopodium (inset from Video 6) in an RGC growth cone imaged with HILO illumination. Acquisition rate: 2 s per frame, playback speed: 12 fps. Red: membrane marker GAP-RFP. Cyan: mNeonGreen-VASP. Scale bar: 1 μm.

**Video 8.** mNeonGreen-ENA localization in filopodia tips during extension (ENA-responding example from Fig. 5C, rotated), in an RGC growth cone imaged with HILO illumination. Acquisition rate: 2 s per frame, playback speed: 12 fps. Red: membrane marker GAP-RFP. Cyan: mNeonGreen-ENA. Scale bar: 1 μm.

**Video 9.** mNeonGreen-ENA localization in filopodia tips during extension (example from Fig. 5F, rotated), in an RGC growth cone imaged with HILO illumination. Acquisition rate: 2 s per frame, playback speed: 12 fps. Red: membrane marker GAPRFP. Cyan: mNeonGreen-ENA. Scale bar: 1 μm.

**Video 10.** Two filopodia tips showing differential response to mNeonGreen-VASP accumulation (examples from Fig. 6A and 6D), in an RGC growth cone imaged with HILO illumination. Acquisition rate: 2 s per frame, playback speed: 12 fps. Red: membrane marker GAP-RFP. Cyan: mNeonGreen-VASP. Scale bar: 1 μm.

